# Behavioural, modeling, and electrophysiological evidence for domain-generality in human metacognition

**DOI:** 10.1101/095950

**Authors:** Nathan Faivre, Elisa Filevich, Guillermo Solovey, Simone Kühn, Olaf Blanke

## Abstract

Metacognition, or the capacity to introspect on one’s own mental states, has been mostly characterized through confidence reports in visual tasks. A pressing question is to what extent the results from visual studies generalize to other domains. Answering this question allows determining whether metacognition operates through shared, domain-general mechanisms, or through idiosyncratic, domain-specific mechanisms. Here, we report three new lines of evidence for decisional and post-decisional mechanisms arguing for the domain-generality of metacognition. First, metacognitive efficiency correlated between auditory, tactile, visual, and audiovisual tasks. Second, confidence in an audiovisual task was best modeled using supramodal formats based on integrated representations of auditory and visual signals. Third, confidence in correct responses involved similar electrophysiological markers for visual and audiovisual tasks that are associated with motor preparation preceding the perceptual judgment. We conclude that the domain-generality of metacognition relies on supramodal confidence estimates and decisional signals that are shared across sensory modalities.

## Introduction

Humans have the capacity to access and report the contents of their own mental states including percepts, emotions, and memories. This capacity, first known as introspection, has been under scrutiny for centuries, and has stirred heated debates among philosophers and psychologists alike (Lyons, 1986). In neuroscience, the reflexive nature of cognition is now the object of research, under the broad scope of the term metacognition (that is, "cognition about cognition”, for reviews see Koriat, 2006; Fleming, Dolan, & Frith, 2012). A widely used method to study metacognition is to have observers do a challenging task (“first-order task”), followed by a confidence judgment regarding their own task performance (“second order task”; Fleming & Lau 2012, see Figure 1 left panel). In this operationalization, metacognitive accuracy can be quantified as the correspondence between subjective confidence judgments and objective task performance. By finely tuning confidence according to performance, a subject with good metacognitive skills will be more confident after correct *vs.* incorrect responses. While some progress has been made regarding the statistical analysis of confidence judgments (Galvin et al., 2003; Maniscalco & Lau, 2012; Barrett, Dienes, & Seth, 2013), and more evidence has been gathered regarding the brain areas involved in metacognitive monitoring (for review see Grimaldi, Lau, & Basso, 2015), the core properties and underlying mechanisms of metacognition remain largely unknown. One of the central questions is whether, and to what extent, metacognitive monitoring should be considered domain-general: is the computation of confidence fully independent of the nature of the task (i.e., domain-generality), or does it also involve task-specific components (i.e., domain-specificity)? According to the domain-generality hypothesis, metacognition would have a quasi-homuncular status, the monitoring of all perceptual processes being operated through a single shared mechanism. Instead, domain-specific metacognition would involve a distributed network of monitoring processes that are specific for each sensory modality or cognitive domain.

The involvement of supramodal, prefrontal brain regions during confidence judgments first suggested that metacognition is partly governed by domain-general rules (e.g., Fleming et al, 2010, 2012; Yokoyama et al., 2010). At the behavioural level, this is supported by the fact that metacognitive performance (Song et al., 2011), and confidence estimates (de Gardelle, Le Corre, & Mamassian, 2014) correlate across subjects between two different visual tasks, as well as between a visual and an auditory task (de Gardelle & Mamassian, 2016). However, the domain-generality of metacognition is challenged by the report of weak or null correlations between metacognitive accuracies across different tasks involving vision, audition, and memory (Ais, Zylberberg, Barttfeld, & Sigman, 2016), and by the finding that functionally co-existing metacognitive brain regions are involved in different tasks, including frontal areas for metaperception and precuneus for metamemory (McCurdy et al., 2013). This anatomo-functional distinction is further supported by the fact that meditation training improves metacognition for memory, but not for vision (Baird, Mrazek, Phillips, & Schooler, 2014). Compared to previous work, the present study sheds new light on the issue of domain-generality by comparing metacognitive monitoring of stimuli from distinct sensory modalities, but during closely-matched first order tasks.

Overall, the evidence supporting the domain-generality of metacognition is mixed, and no mechanism explaining its hypothetical origins has been proposed. Two non-mutually exclusive mechanisms responsible for domain-generality can be considered. First, metacognition may be domain-general in case monitoring operates on supramodal confidence estimates, computed with an identical format or neural code across different tasks or sensory modalities (mechanism 1; Pouget, Drugowitsch, & Kepecs, 2016). Second, metacognition may be domain-general in case a non-perceptual signal drives the computation of confidence estimates (mechanism 2). Among them, likely candidates are decisional cues such as reaction times during the first-order task, as they are present no matter the sensory modality at play, and are thought to play an important role for confidence estimates (Yeung & Summerfield, 2012).

Here we sought to test the hypothesis of domain-generality in metacognition, and directly explore the two above-mentioned mechanisms. At the behavioral level, we first investigated the commonalities and specificities of metacognition across sensory domains including touch, a sensory modality that has been neglected so far. Namely, we examined correlations between metacognitive performance during a visual, auditory, and tactile discrimination task (Experiment 1). Next, extending our paradigm to conditions of audiovisual stimulation, we quantified for the first time the links between unimodal and multimodal metacognition (Deroy et al., 2016), and assessed through computational modeling how multimodal confidence estimates are built (Experiment 2). This allowed us to assess if metacognition is domain-general because of the generic format of confidence. Finally, we investigated the neural mechanisms of unimodal and multimodal metacognition and repeated Experiment 2 while recording 64-channel electroencephalography (EEG, Experiment 3). This allowed us to identify neural markers with high temporal resolution, focusing on those preceding the response in the first-order task to assess if metacognition is domain-general because of the presence of decisional cues (mechanism 2; Boldt et al., 2015). Namely, we assessed the presence of domain-general versus domain-specific mechanisms of metacognition during the motor preparation preceding the perceptual first-order task, as quantified by event-related potentials (ERPs) and alpha suppression over sensorimotor regions. The present data reveal (1) correlations in metacognitive behavioral efficiencies across different unimodal and bimodal perception, (2) computational evidence for integrative, supramodal representations during audiovisual confidence estimates (mechanism 1), and (3) the presence of similar neural markers of domain-general metacognition preceding the first-order task (mechanism 2). Altogether, these behavioural, computational, and neural findings provide non-mutually exclusive mechanisms explaining the domain-generality of metacognition during human perception.

**Figure 1:**
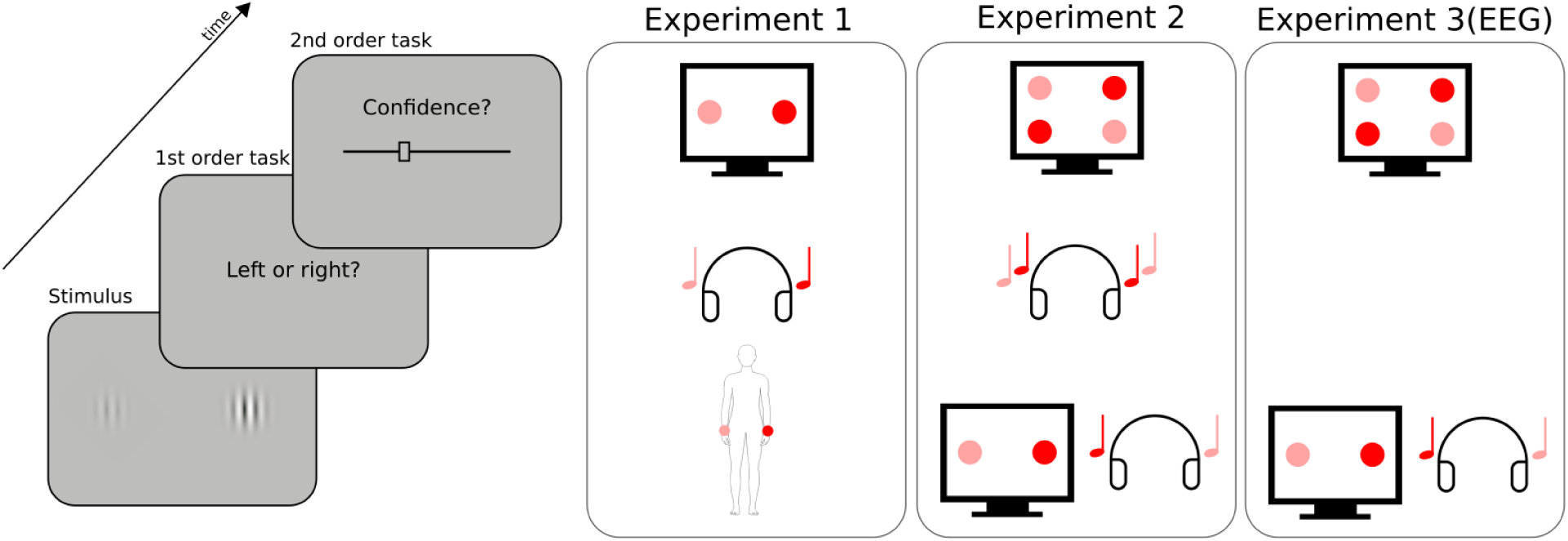
Experimental procedure. Participants had to perform a perceptual task on a stimulus (first order task), and then indicate their confidence in their response by placing a cursor on a visual analog scale (second order task). The types of stimuli and first order task varied across conditions and experiments, as represented schematically on the right panel. In Experiment 1, a pair of two images, sounds, or tactile vibrations was presented on each trial. The stimuli of each pair were lateralized and differed in intensity (here high intensity is depicted in red, low intensity in pink). The first order task was to indicate whether the most intense stimulus was located on the right (as depicted here) or left side. In Experiment 2, either two pairs of two images (unimodal visual condition), two sounds (unimodal auditory condition), or one pair of two images with one pair of two sounds (bimodal audiovisual condition) were presented on each trial. The first-order task was to indicate whether the most intense stimulus of each pair were both on the same side (congruent trial), or each on a different side (incongruent trial, as depicted here). Experiment 3 was a replication of Experiment 2 including EEG recordings, focusing on the unimodal visual condition and the bimodal audiovisual condition. The order of conditions within each experiment was counterbalanced across participants.

## Results

### Experiment 1

We first aimed at comparing metacognitive accuracy across the visual, auditory, and tactile modalities. Participants were presented with a pair of simultaneous stimuli at a right and left location, and asked to indicate which of the two stimuli had the highest intensity (Figure 1 right panel). In this way, the first-order task consisted in a 2-alternative forced choice on visual, auditory, or tactile intensity (i.e., respectively contrast, loudness, or force). After each choice, participants reported their confidence on their previous response (second-order task) (Figure 1 left panel). The main goal of this experiment was to test the existence of correlations in metacognitive efficiency between sensory modalities. Before examining them, we report general results of type 1 and type 2 performances. First, we aimed to equate first-order performance in the three modalities using a 1-up/2-down staircase procedure (Levitt, 1971). Although this approach prevented large inter-individual variations, some minor differences across modalities subsisted, as revealed by a one-way ANOVA on d’ measuring first order sensitivity [F(1.92,26.90) = 8.76, p < 0.001, η_p_^2^ = 0.38] (Figure 2a). First-order sensitivity was lower in the auditory condition [mean d’ = 1.20 ± 0.05 (95% CI)] as compared to the tactile [mean d’ = 1.37 ± 0.07, p = 0.002] and visual conditions [mean d’ = 1.33 ± 0.07, p = 0.004] (Figure 2a; see SI for further analyses). This aspect of our results is likely due to the difficulty of setting perceptual thresholds with adaptive staircase procedures. Importantly however, it does not prevent us from comparing metacognitive performance across senses, as the metrics of metacognitive performance we used are independent of first-order sensitivity. Indeed, metacognitive sensitivity was estimated with meta-d’, a response-bias free measure of how well confidence estimates track performance on the first-order task (Maniscalco & Lau, 2012). A one-way ANOVA on meta-d’ revealed a main effect of condition [F(1.93,25.60) = 5.92, p = 0.009, η_p_^2^ = 0.30] (Figure 2b). To further explore this main effect and rule out the possibility that it stemmed from differences at the first-order level, we normalized metacognitive sensitivity by first-order sensitivity (i.e., meta-d’/d’), to obtain a pure index of metacognitive performance called metacognitive efficiency. Only a trend for a main effect of condition was found [F(1.76,24.61) = 3.16, p = 0.07, η_p_^2^ = 0.18] (Figure 2c), revealing higher metacognitive efficiency in the visual [mean ratio = 0.78 ± 0.13] *vs.* auditory domain [mean meta-d’/d’ ratio = 0.61 ± 0.15; paired t-test: p = 0.049]. The difference in metacognitive efficiency between the visual and the tactile conditions [mean ratio = 0.70 ± 0.10] did not reach significance [paired t-test: p = 0.16]. These results and others that are secondary to test the domain-generality hypothesis of metacognition are further discussed in SI.

**Figure 2:**
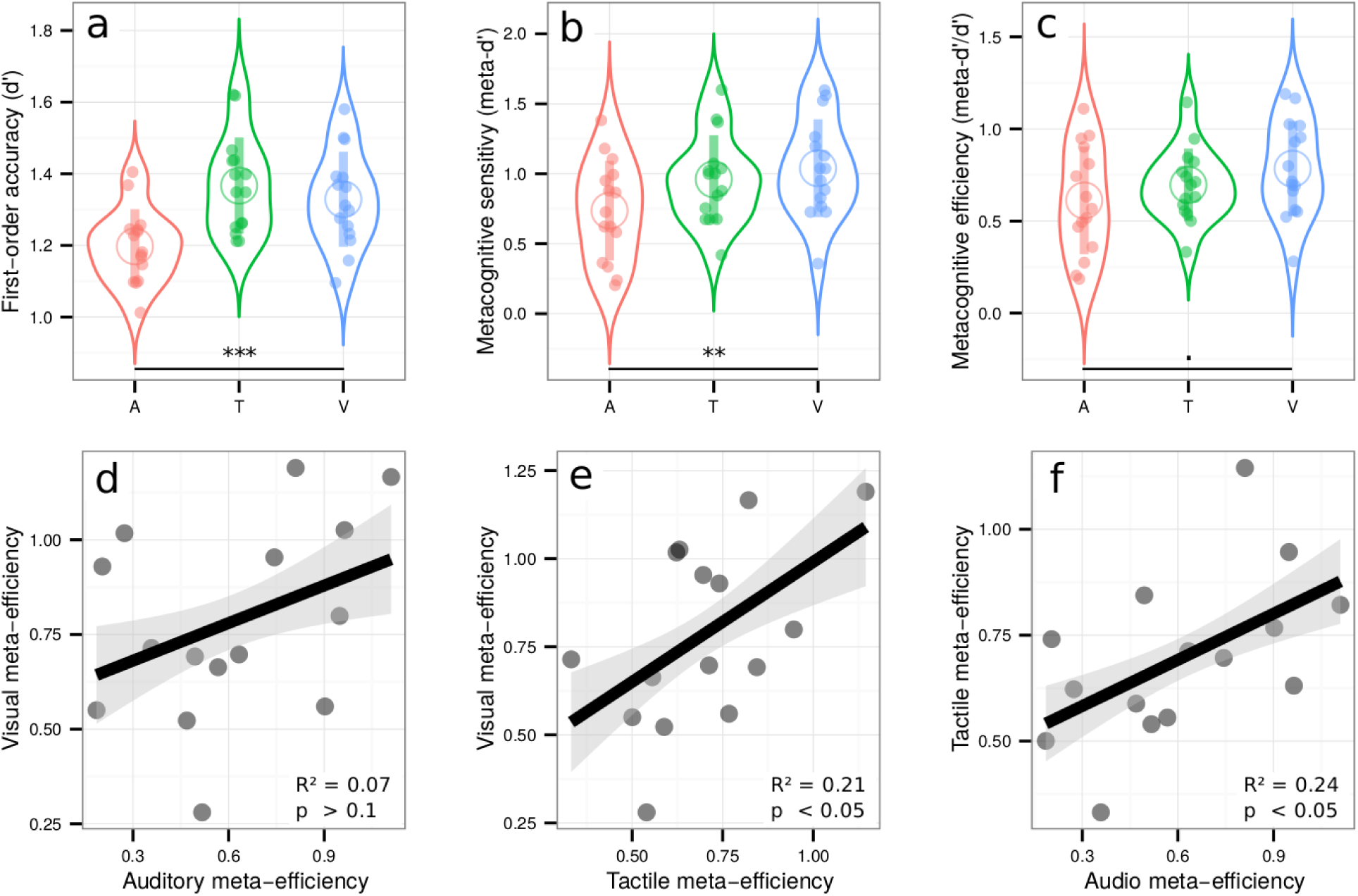
Upper row: Violin plots representing first order sensitivity (a: d’), metacognitive sensitivity (b: meta-d’), and metacognitive efficiency (c: meta-d’/d’) in the auditory (A, in red), tactile (T, in green), and visual modalities (V, in blue). Full dots represent individual data points. Empty circles represent average estimates. Error bars represent the standard deviation. The results show that independently of first-order performance, metacognitive efficiency is better in vision compared to audition. **Lower row**: correlations between metacognitive efficiencies in the visual and auditory conditions (3d), visual and tactile conditions (3e), and tactile and auditory conditions (3f). The results show that metacognitive efficiency correlates across sensory modalities, in favor of the domain-general hypothesis. *** p < 0.001, ** p < 0.01, + p < 0.1.

Crucially, we found positive correlations between metacognitive efficiency in the visual and tactile conditions [adjusted R^2^ = 0.21, p = 0.047] (Figure 2e), and in the auditory and tactile conditions [adjusted R2 = 0.24, p = 0.038] (Figure 2f). (The data were inconclusive regarding the correlation between the visual and auditory condition [adjusted R^2^ = 0.07, p = 0.17, Bayes Factor = 0.86] (Figure 2d)). These results reveal shared variance between auditory, tactile, and visual metacognition, in line with the domain-generality hypothesis. Moreover, the absence of any correlation between first-order sensitivity and metacognitive efficiency in any of the conditions [all adjusted R^2^ < 0; all p-values > 0.19], rules out the possibility that such domain-generality during the second-order task was confounded with first-order performance.

### Experiment 2

Experiment 1 revealed correlational evidence for the domain-generality of perceptual metacognition across three modalities. A previous study (McCurdy et al., 2013), however, dissociated brain activity related to metacognitive accuracy in vision versus memory, despite clear correlations at the behavioral level. Thus, correlations between modalities are compelling, but not sufficient to support the domain-generality hypothesis. We therefore put the evidence of experiment 1 to a stricter test in Experiment 2, by comparing metacognitive efficiency for unimodal vs. bimodal, audiovisual stimuli. We reasoned that if metacognitive monitoring operates independently from the nature of sensory signals from which confidence is inferred, confidence estimates should be as accurate when made on unimodal or bimodal signals. In contrast, if metacognition operated separately in each sensory modality, one would expect that metacognitive efficiency for bimodal stimuli would only be as high as the minimal metacognitive efficiency for unimodal stimuli. Besides mere comparisons, the domain-generality hypothesis also implies the existence of correlations between unimodal and bimodal metacognitive efficiencies, thereby extending the correlations across distinct unimodal stimulations reported in Experiment 1 to bimodal stimulations. Participants performed three different perceptual tasks, all consisting in a congruency judgment between two pairs of stimuli (Figure 1, right panel). In the unimodal visual condition, participants indicated whether the most contrasted stimuli of each pair were situated on the same or different side of the screen. In the unimodal auditory condition, they indicated whether the loudest sounds of each pair were played in the same ear or in two different ears. In the bimodal audiovisual condition, participants indicated whether the side corresponding to the most contrasted Gabor patch of the visual pair corresponded with the side of the loudest sound of the auditory pair. The staircase procedure minimized variations in first-order sensitivity [F(1.75,22.80) = 2.12, p = 0.15, ηρ^2^ = 0.14], such that sensitivity in the auditory [mean d’ = 1.31 ± 0.12], audiovisual [mean d’ = 1.38 ± 0.12], and visual conditions [mean d’ = 1.25 ± 0.11] were similar (Figure 3a, and SI for further analyses). We found a main effect of condition for both metacognitive sensitivity [meta-d’: F(1.98,25.79) = 4.67, p = 0.02, ηρ^2^ = 0.26], and metacognitive efficiency [ratio meta-d’/d’: F(1.95,25.40) = 6.63, p = 0.005, ηp^2^ = 0.34] (Figure 3b and 3c, respectively). Pairwise comparisons revealed higher metacognitive efficiency in the visual [mean ratio = 0.94 ± 0.19] vs. auditory [mean meta-d’/d’ ratio = 0.65 ± 0.17; paired t-test: p = 0.005] and audiovisual domains [mean meta-d’/d’ ratio = 0.70 ± 0.15; paired t-test: p = 0.02]. As auditory and audiovisual metacognitive efficiencies were not different [p = 0.5, BF = 0.38], the differences in metacognitive efficiency are likely to stem from differences between auditory and visual metacognition, as found in Experiment 1. Importantly, congruency judgments required that participants responded on the basis of the two presented modalities. Thus, the fact that metacognitive efficiency is similar in the audiovisual and auditory tasks implies that the resolution of confidence estimates in the bimodal condition is as good as that in the more difficult unimodal condition (in this case, auditory), despite it requiring the analysis of two sources of information.

**Figure 3:**
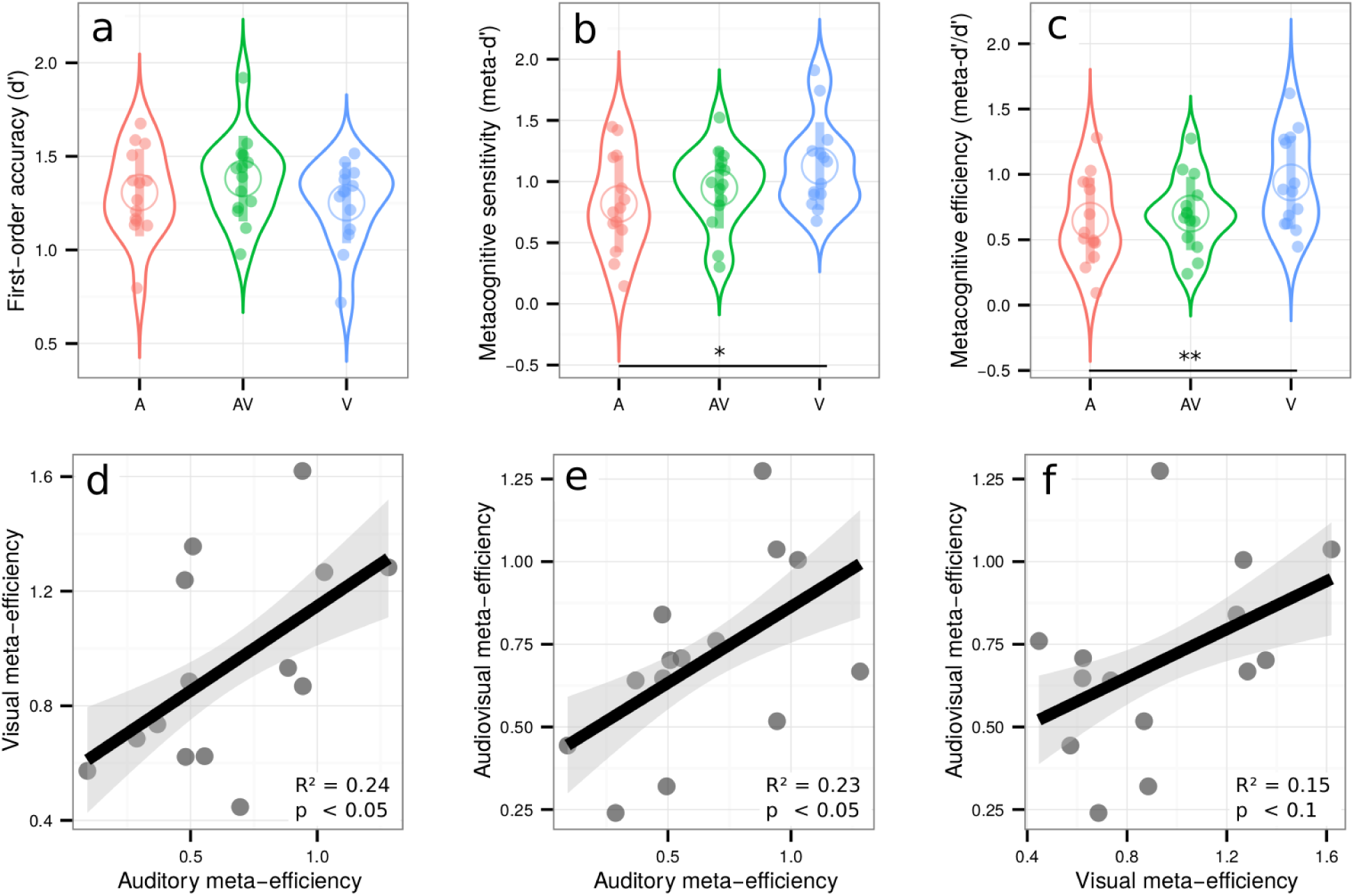
Upper row: Violin plots representing first order sensitivity (3a: d’), metacognitive sensitivity (3b: meta-d’), and metacognitive efficiency (3c: meta-d’/d’) in the auditory (A, in red), audiovisual (AV, in green), and visual modalities (V in blue). Full dots represent individual data points. Empty circles represent average estimates. Error bars represent the standard deviation. The results show that independently of first-order performance, metacognitive efficiency is better for visual stimuli vs. auditory or audiovisual stimuli, but not poorer for audiovisual vs. auditory stimuli. Lower row: correlations between metacognitive efficiency between conditions. **Lower row**: correlations between relative metacognitive efficiency in the visual and auditory conditions (3d), audiovisual and auditory conditions (3e) and audiovisual and visual conditions (3f). The results show that metacognitive efficiency correlates between unimodal and bimodal perceptual tasks, in favor of the domain-general hypothesis. ** p < 0.01, * p < 0.05.

Crucially, we found again correlations between metacognitive efficiency in the auditory and visual conditions [adjusted R^2^ = 0.24, p = 0.043] (Figure 3d), as in experiment 1; more importantly, we also found correlations between metacognitive efficiency in the auditory and audiovisual conditions [adjusted R^2^ = 0.23, p = 0.046] (Figure 3e) and a trend between metacognitive efficiency in the visual and audiovisual conditions [adjusted R^2^ = 0.15, p = 0.097] (Figure 3f). This contrasted with no correlations between first-order sensitivity and metacognitive efficiency in any of the conditions [all R^2^ < 0.06; all p > 0.19] except in the visual condition, where high d’ was predictive of low meta-d’/d’ values [R^2^ = 0.39, p = 0.01]. The absence of such correlations in most conditions makes it unlikely that relations in metacognitive efficiency were driven by similarities in terms of first-order performance. In addition to the equivalence between the resolution of unimodal and bimodal confidence estimates, the correlations in metacognitive efficiency between unimodal and bimodal conditions suggest that metacognitive monitoring for unimodal vs. bimodal signals involves shared mechanisms (i.e. domain-generality).

### Computational models of confidence estimates for bimodal signals

Using the data from experiment 2, we next sought to reveal potential mechanisms underlying the computation of confidence in the bimodal condition. For this, we first modeled the proportion of trials corresponding to high vs. low confidence in correct vs. incorrect type 1 responses, in the unimodal auditory and unimodal visual conditions separately. Each condition was represented by a 2-dimensional signal detection theory (SDT) model with standard assumptions and only 2 free parameters per participant, namely internal noise σ and confidence criterion *c* (see Figure 4, Methods and SI for details). This simple model accounted for more than half the total variance in participants’ proportion of responses both in the unimodal visual [R^2^ = 0.68] and unimodal auditory conditions [R^2^ = 0.57]. We then combined the fitted parameter values under different rules to estimate and compare their fits to the audiovisual data. Note that with this procedure, and unlike the fits to the unimodal conditions, the data used to estimate the model parameters were different from those on which the model fits were compared. We evaluated different models systematically by grouping them into three families, varying in degree of domain-generality. We present here the best model from each family (see figure 4 and figure S5 for schematics of all models tested). The *domain-general model* echoes the unimodal models and represents the highest degree of integration: here, confidence is computed on the basis of the joint distribution of the auditory and visual modalities. The *comparative model* assumes that confidence is computed separately for each modality and in a second step combined into a single summary measure. The *single-modality model* assumes that confidence varies with the internal signal strength of a single modality and therefore supposes no integration of information at the second-order level. We compared these different models by calculating their respective BIC weights (BICw, Burnham & Anderson, 2002; Solovey, Graney, & Lau, 2014), which quantify the relative evidence in favour of a model in relation to all other models considered.

**Figure 4:**
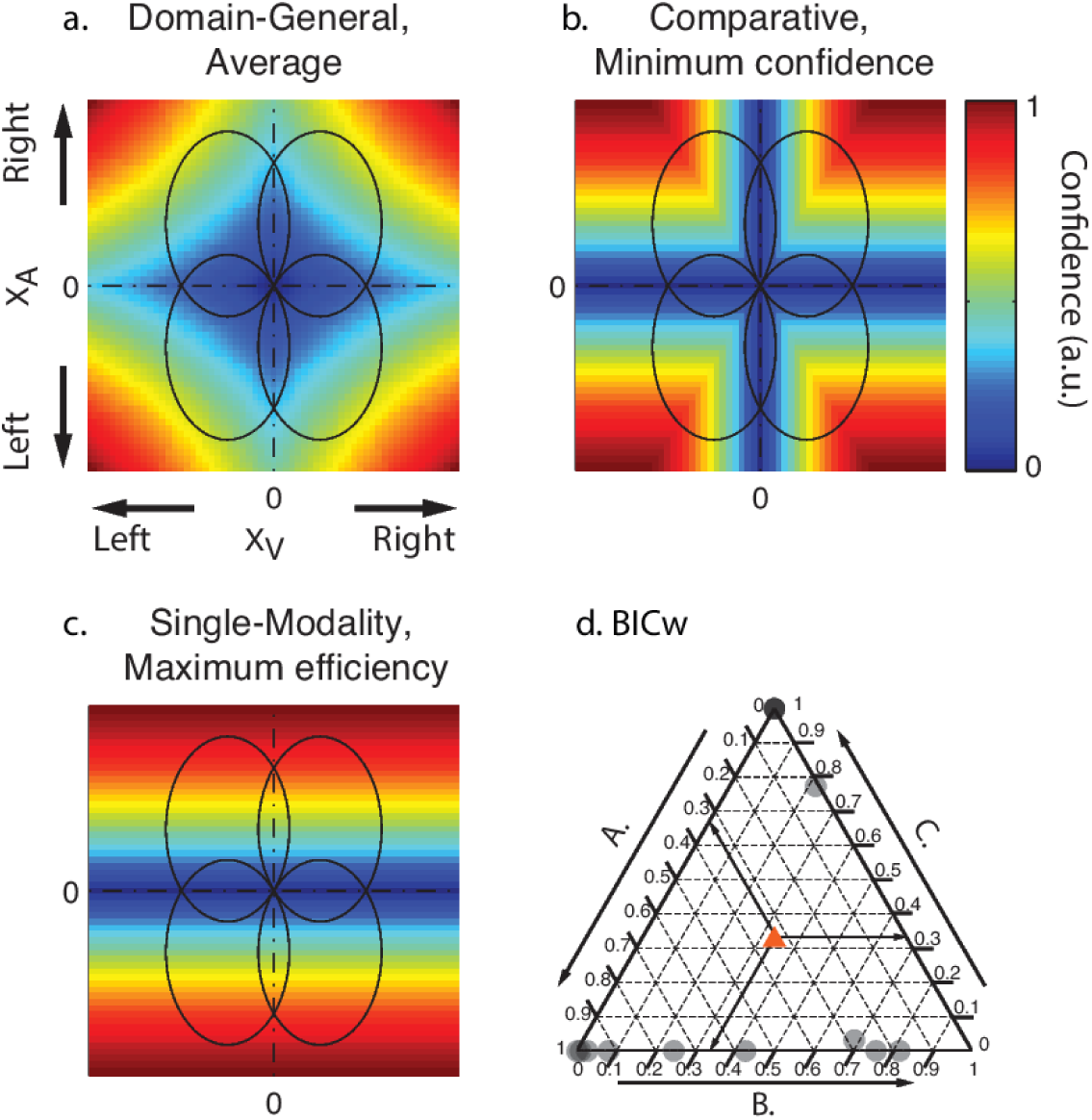
Model predictions (a, b, c): Ellipses represent the partially overlapping bivariate internal signal distributions for each of the stimulus combinations, represented at a fixed density contour. The top right quadrant corresponds to congruent stimuli, where the stimuli in each pair were stronger on the right side. The colours represent the predicted confidence, normalized to the interval [0,1] for every combination of internal signal strength for each stimulus pair (Xa, Xv). Note that for models A and B, confidence increases with increasing internal signal level in both modalities, whereas in the single-modality model C, confidence depends on the signal strength of only one modality. **(d.) Individual BIC weights for the three model fits in the audiovisual condition.** The arrows show how to read the plot from an arbitrary data point in the diagram, indicated with a red triangle. Consider that the sum of the BICw for all models (A., B. and C.) amounts to 1 for each participant. To estimate the relative BICw of each model for any given participant, take the lines parallel to the vertex labeled 1 for that model. The intersection between the line parallel to the vertex and the triangle edge corresponding to the model indicates the BICw.

By examining individual BICw in a ternary plot (Figure 4d; see also figure S4 for individual model fits), we found that the best model for most participants was either the *domain-general* or the *comparative model*, whereas the BICw for the *single-modality model* was equal to 0. Yet, we note that the *single-modality* model is also plausible, as it does predict the responses of four participants better than any of the other two models. Taken together these computational results suggest that most participants computed confidence in the bimodal task by using information from the two modalities under a supramodal format that is independent of the sensory modality, in agreement with the first mechanism for domain-general metacognition we introduced. We conclude that the confidence reports for audiovisual signals arise either from the joint distribution of the auditory and visual signals (*domain-general model*), or are computed separately for distinct modalities, and then combined into a single supramodal summary statistic (*comparative model*). These two models are therefore good candidates to explain the domain-generality of metacognition. Besides this first mechanism in favor of the domain-general hypothesis, we next sought to assess if metacognition was domain-general due to the influence of decisional cues that are shared between sensory modalities (see introduction: mechanism 2).

Our modeling results suggest that confidence estimates are encoded in a supramodal format, compatible with the domain-generality hypothesis for metacognition. Notably however, apparent domain-generality in metacognition could arise in case non-perceptual signals are taken as inputs for the computation of confidence. In models implying a decisional locus for metacognition (Yeung & Summerfield, 2012), stimulus-independent cues such as reaction times during the first-order task take part in the computation of confidence estimates. This is empirically supported by a recent study showing that confidence in correct responses is decreased in case response-specific representations encoded in the premotor cortex are disrupted by transcranial magnetic stimulation (Fleming et al., 2015). In the present study, decisional parameters were shared across sensory modalities, since participants used a keyboard with their left hand to perform the first-order task for all sensory modalities. To extend our modeling results and assess whether domain-generality in metacognition also involves a decisional locus (mechanism 2 discussed above), we examined how participants used their reaction times to infer confidence in different conditions. Specifically, we quantified the overlap of first-order reaction times distributions corresponding to correct vs. incorrect responses, as a summary statistic representing how reaction times differ between correct and incorrect trials. We measured how reaction time overlap correlated with the overlap of confidence ratings after correct vs. incorrect first-order responses, which is a summary statistic analogous to ROC-based methods typically used to quantify metacognitive sensitivity with discrete confidence scales (Fleming & Lau, 2012). If confidence involves a decisional-locus, one would expect a correlation between confidence overlap and reaction time overlap, so that participants with the smallest confidence overlap (i.e., highest metacognitive sensitivity) are the ones with the smallest reaction times overlap (i.e., distinct reaction times in correct vs. incorrect responses). Interestingly in Experiment 1 the correlation strength mirrored the difference in metacognitive efficiency we found between sensory modalities: higher correlations were found in the visual domain (adjusted R^2^ = 0.54, p = 0.002; average metacognitive efficiency = 0.78 ± 0.13), compared to the tactile (adjusted R^2^ = 0.26, p = 0.03; average metacognitive efficiency = 0.70 ± 0.10) and auditory domains (adjusted R^2^ = -0.06, p = 0.70; average metacognitive efficiency = 0.61 ± 0.15; see figure S1). This suggests that decisional parameters such as reaction times in correct vs. incorrect trials may inform metacognitive monitoring, and may be used differently depending on the sensory modality with a bigger role in visual than in tactile and auditory tasks. Importantly, even though such correlations between reaction time overlap and confidence overlap would be expected in experiments containing a mixture of very easy and very difficult trials, the correlations in the visual and tactile modalities reported above persisted even after the variance of perceptual evidence was taken into account using multiple regressions. This rules out the possibility that these correlations are explained by variance in task difficulty. This pattern of results was not found in Experiment 2 (i.e. no correlation between reaction times and confidence overlaps; all R^2^ < 0.16, all p > 0.1), but replicated in Experiment 3 as further detailed below.

### Experiment 3.

The aim of experiment 3 was three-fold. First and foremost, we sought for the first time to document the potential common and distinct neural mechanisms underlying unimodal and bimodal metacognition. Following the link between reaction times and metacognitive efficiency uncovered in Experiment 1, we expected to find domain-general neural markers of metacognition preceding the first-order task, as quantified by the amplitude of event-related potentials (ERPs) as well as in alpha suppression over the sensorimotor cortex prior to key press (Pfurtscheller & Da Silva, 1999). Second, we aimed at replicating the behavioural results from Experiment 2, especially the correlation between visual and audiovisual metacognitive efficiency. Third, we aimed at estimating the correlations between confidence and reaction times overlap on a new group of participants. Therefore, we tested participants on these two conditions only.

### Behavioural data

The staircase procedure minimized variations in first-order sensitivity [t(17) = 0.3, p = 0.76, d = 0.07], such that sensitivity in the audiovisual [mean d’ = 1.15 ± 0.07] and visual conditions [mean d’ = 1.17 ± 0.05] were similar (see SI for further analyses). No difference in metacognitive sensitivity was found between conditions [t(17) = 0.78, p = 0.44, d = 0.09] or efficiency [t( 17) = 0.78, p = 0.44, d = 0.08]. Crucially, we replicated our main results from Experiment 2, as we found a positive significant correlation between relative metacognitive accuracy in the audiovisual and visual conditions [adjusted R^2^ = 0.47, p < 0.001], and no correlation between first-order sensitivity and metacognitive efficiency in either condition [both R^2^ < 0.01; both p-values > 0.3] (Figure 5). Regarding the decisional locus of metacognition, Experiment 3 confirmed the results of Experiment 1: reaction time and confidence overlaps correlated more in the visual condition (adjusted R^2^ = 0.41, p = 0.003), than in the audiovisual condition (adjusted R^2^ = -0.05, p = 0.70), suggesting that decisional parameters such as reaction times may inform metacognitive monitoring, although differently between the visual and audiovisual conditions. Altogether, these behavioral results from three experiments with different subject samples confirm the existence of shared variance in metacognitive efficiency between unimodal and bimodal conditions, and do not support major group differences between them. Further, they support the role of decisional factors such as reaction times estimates, as predicted when considering a decisional locus for metacognition.

**Figure 5:**
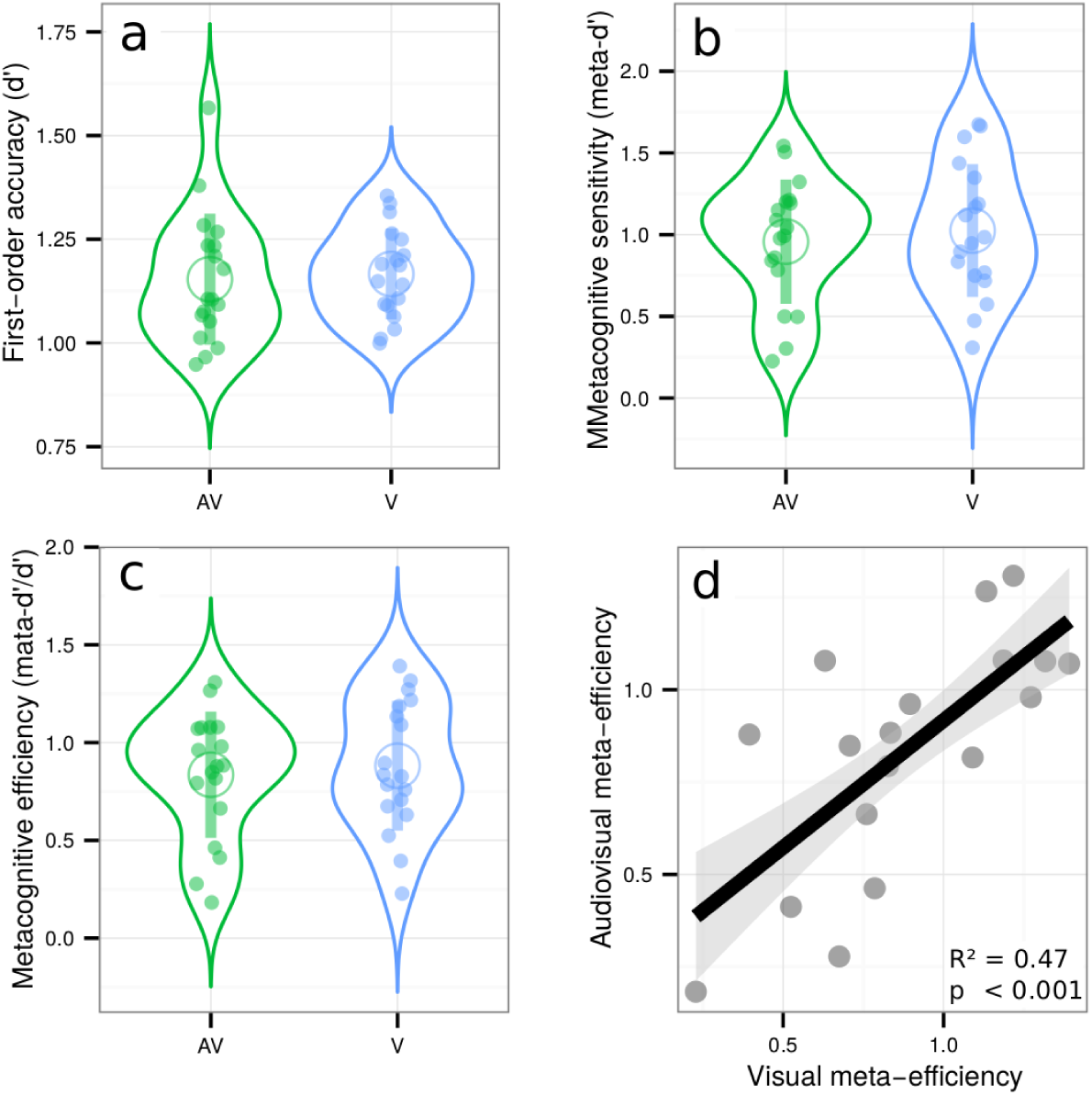
Violin plots represent first-order sensitivity (5a: d’), metacognitive sensitivity (5b: meta-d’), and metacognitive efficiency (5c: meta-d’/d’) in the audiovisual (AV, in green), and visual conditions (V, in blue). Full dots represent individual data points. Empty circles represent average estimates. Error bars represent the standard deviation. The results show no difference between visual and audiovisual metacognitive efficiency. 5d represents the correlation between metacognitive efficiency in the audiovisual and visual conditions.

### EEG data

Next, we explored the neural bases of visual and audiovisual metacognition, focusing on the decisional locus of confidence by measuring ERPs locked to the type 1 response, taking into account the differences in type 1 reaction times between the visual and audiovisual tasks (562ms shorter in the visual condition on average: t(17) = 6.30, p < 0.001). Since we showed that decisional parameters such as reaction times inform metacognitive monitoring, this analysis was carried out on a set of scalp electrodes over the right sensorimotor cortex that included the left hand representation with which participants performed the first-order task (see Boldt & Yeung for findings showing that parietal scalp regions also correlate with confidence prior to response). Incorrect type 1 responses were not analyzed as the lower-bound of the confidence scale we used corresponded to a “pure guess”, and therefore did not allow disentangling detected vs. undetected errors.

We first compared the average ERP amplitude time-locked to the onset of correct type 1 responses as a function of confidence, using linear mixed models with condition as a fixed effect (visual vs. audiovisual, see Methods and SI for details). This analysis revealed main effects, whereby the modulations of ERP amplitudes by confidence were similar in the visual and audiovisual condition, and interaction effects, whereby the amplitude modulations differed across conditions. A first main effect of confidence was found early before the type 1 response, underlying a negative relationship between ERP amplitude and confidence (-600 to -550 ms; p < 0.05, fdr-corrected, see figure 6a, left panel, showing the grand average between the visual and audiovisual condition). A second main effect of confidence peaked at -300 ms (-400 to -100 ms; p < 0.05, fdr-corrected) so that trials with high confidence reached maximal amplitude 300 ms before key press. These two effects are characterized by an inversion of polarity from an early-negative to a late-positive relationship, which has been linked to selective response activation processes (i.e., lateralized readiness potentials, see Eimer, 1998 for review, and Bujan et al., 2009 for previous results in metamemory). Thus, the present data show that sensorimotor ERP also contribute to metacognition as they seemed impacted by confidence both in the audiovisual and visual conditions. Of note, confidence modulated the amplitude and not the onset latency of the ERP, which suggests that the timing of response selection itself does not depend on confidence. We complemented this ROI analysis by exploring the relation between confidence and ERP amplitude for all recorded electrodes (figure 6a, right panel). This revealed that the later effect 300ms before key press was centered on centro-parietal regions (i.e., including our region of interest; p < 0.001) as well as more frontal electrodes, potentially in line with several fMRI studies reporting the role of the prefrontal cortex for metacognition (Fleming et al., 2010; McCurdy et al., 2013, see Grimaldi et al., 2015 for review). The linear mixed model analysis also revealed significant interactions, indicating that the modulation of ERP amplitude as a function of confidence was significantly stronger in the visual condition, with again one early (−750 to −600ms) and late component (−350 to −150ms; Figure 6b left panel)). Topographical analysis of these interactions implicated frontal and parieto-occipital electrodes. These results at the neural level are consistent with our behavioural data, since we found that reaction times have more influence on the computation of confidence in the visual compared to the audiovisual condition.

**Figure 6:**
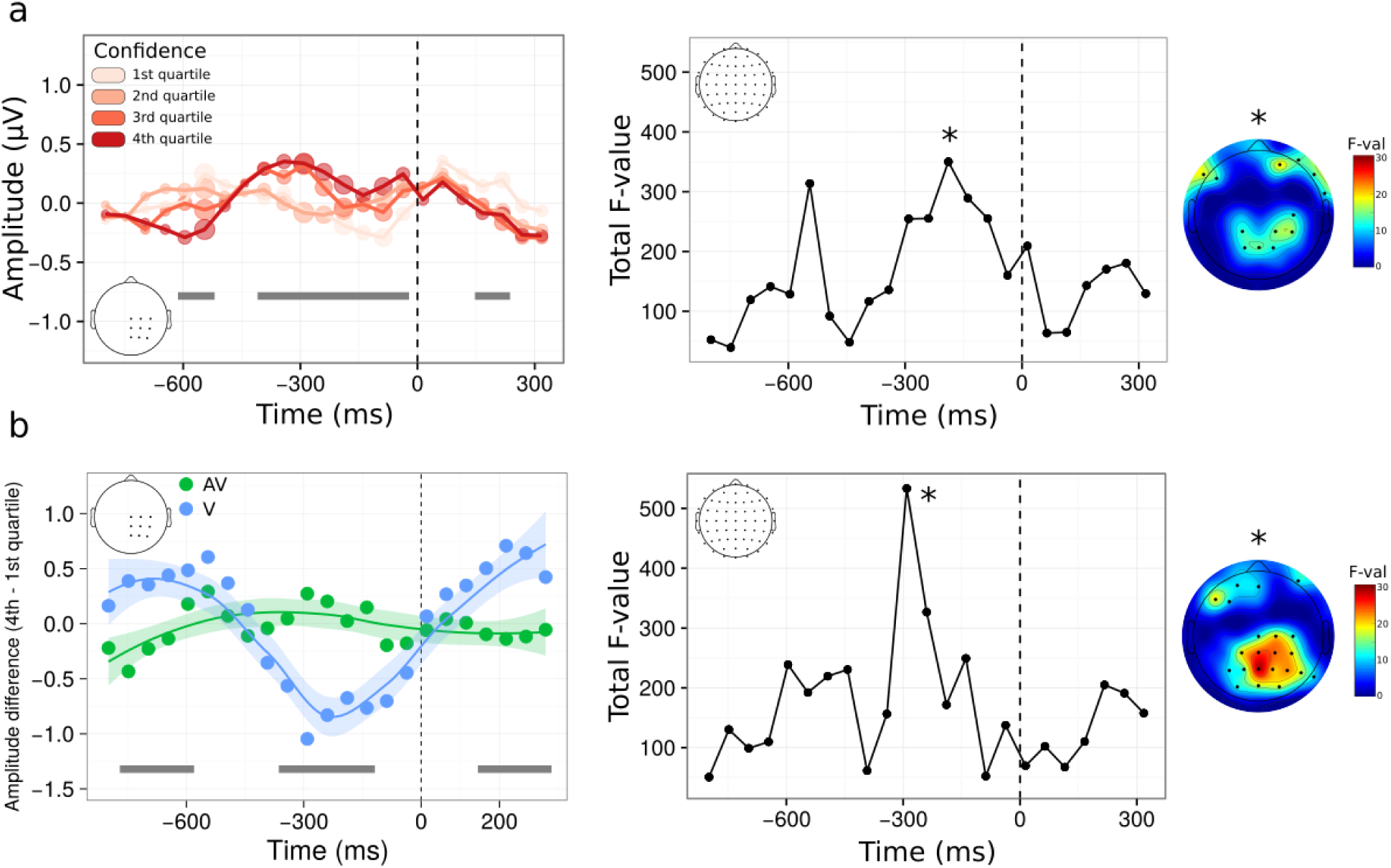
Voltage amplitude time-locked to correct type 1 responses as a function of confidence. **a.** left panel: time course of the main effect of confidence within a pre-defined ROI. Although raw confidence ratings were used for the statistical analysis, they are depicted here as binned into four quartiles, from quartile 1 corresponding to trials with the 25% lowest confidence ratings (light pink), to quartile 4 corresponding to trials with the 25% highest confidence ratings (dark red). **Right panel**: same analysis as shown in (a) on the whole scalp. The plot represents the time-course of the summed F-value over 64 electrodes for the main effect of confidence. The topography where a maximum F-value is reached (*) is shown next to each plot. **b. left panel**: time course of the interaction between confidence and condition following a linear mixed model analysis within the same ROI as in (a). Although raw confidence ratings were used for the statistical analysis, the plot represents the difference in voltage amplitude between trials in the 4^th^ vs. 1^st^ confidence quartile. **Right panel**: same analysis as shown in (b) on the whole scalp, with corresponding topography. In all plots, grey bars correspond to significant main effects (a) or interactions (b), with p < 0.05 fdr-corrected. Significant effects on topographies are highlighted with black stars (p < 0.001, uncorrected).

Complementary to ERP amplitude, we also analyzed oscillatory alpha power as a signature of motor preparation (i.e, pre-movement related desynchronization, see Pfurtscheller & Da Silva, 1999). Results of the linear mixed model analysis revealed a sustained main effect of confidence starting 300ms before key press and continuing until 200ms after the type 1 response (p < 0.05 fdr-corrected), showing a negative relationship between confidence and alpha power (i.e., alpha suppression, figure 7a, left panel). Note that, opposite to what we found in the amplitude domain, the main effect of confidence on alpha power was found even after a first-order response was provided. Likewise, the topographical analysis revealed a different anatomical localization than the effect we found in the amplitude domain, with more posterior, parieto-occipital electrodes involved. This suggests that alpha suppression prior to type 1 response varies as a function of confidence non-differentially in both the audiovisual and visual conditions. The linear mixed model analysis also revealed a main effect of condition, with higher alpha power in the visual vs. audiovisual condition (figure 7b, left panel). This could be related to the fact that the audiovisual task was judged more demanding by participants, as reflected by their longer type 1 reaction times. Finally, significant interactions between confidence and condition were found, with topographical locations predominantly within frontal electrodes. Taken together, the main effects of confidence on voltage amplitude and alpha power reveal some of the markers validating the domain-generality hypothesis at a decisional locus. These are likely to be part of a bigger set of neural mechanisms, operating at a decisional, but also post-decisional locus that was not explored here (Pleskak & Busemeyer, 2010). The existence of significant interactions reveals that some domain-specific mechanisms are also at play during metacognition, which accounts for the unexplained variance when correlating metacognitive efficiencies across modalities at the behavioral level.

**Figure 7:**
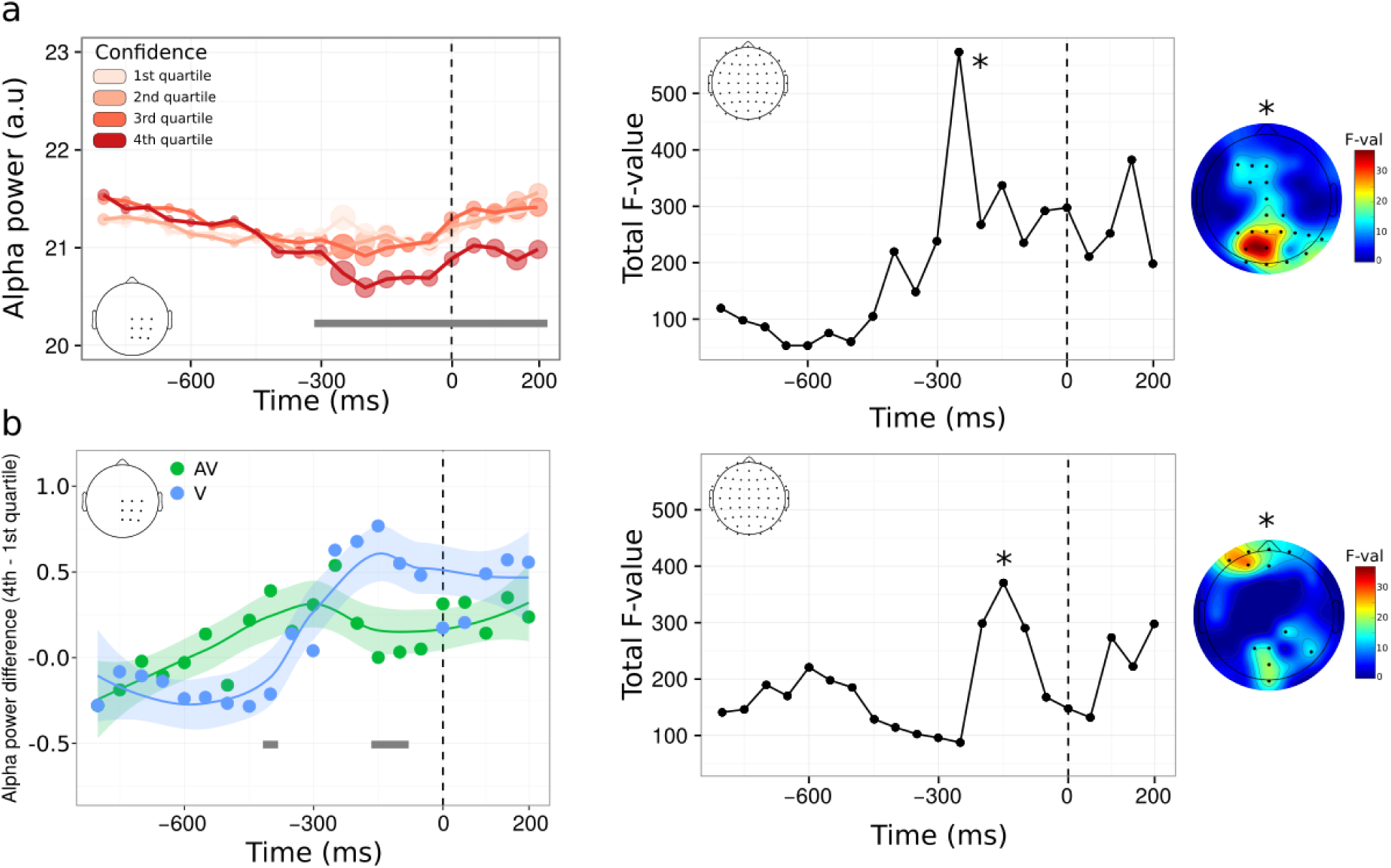
Alpha power time-locked to correct type 1 responses as a function of confidence. The legend is identical to that of figure 6, with alpha power instead of voltage amplitude.

## Discussion

Is perceptual metacognition domain-general, with a common mechanism for distinct tasks and sensory modalities, or is it domain-specific, with idiosyncratic mechanisms for each task and sensory modality? As of today, this issue remains unsettled because the vast majority of experiments on metacognitive perception only involved the visual modality (but see Ais, 2016; de Gardelle 2016). In vision, Song and colleagues (2011) found that about half of the variance in metacognitive sensitivity during a contrast discrimination task was explained by metacognitive sensitivity in an orientation discrimination task, suggesting some level of generality across different tasks within the visual domain. Likewise, roughly a quarter of the variance in metacognitive sensitivity during a contrast discrimination task was explained by metacognitive sensitivity during a memory task involving words presented visually (McCurdy et al., 2013). Here, we extend these studies beyond vision with several new experiments aimed at assessing the generality of metacognition across three sensory modalities as well as conjunctions of two sensory modalities. In Experiment 1 we tested participants in three different conditions, which respectively required discriminating the side on which visual, auditory or tactile stimuli were most salient. We found positive correlations between metacognitive efficiency across sensory modalities, and ruled out the possibility that these correlations stemmed from differences in firstorder performances (Maniscalco & Lau, 2012). These results extend the report by Ais and colleagues (2016) and de Gardelle and colleagues (2016) of similarities between auditory and visual metacognition to auditory, tactile, and visual laterality discrimination tasks, and therefore support the existence of a common mechanism underlying metacognitive judgments in three distinct sensory modalities.

In Experiment 2, we further extended these results to a different task and also generalized them to bimodal stimuli (Deroy et al., 2016). First, using a first-order task that required congruency rather than laterality judgments, we found again that metacognitive efficiency in an auditory task correlated with metacognitive efficiency in a visual task. Second, we designed a new condition in which participants had to perform congruency judgments on bimodal, audiovisual, signals, which required the information from both modalities to be taken into account. Three further observations from these conditions support the notion of domain-generality in perceptual metacognition. First, we observed that metacognitive efficiency in the audiovisual task was indistinguishable from that in the unimodal auditory task, suggesting that the computation of joint confidence is not only possible but can also occur at no behavioral additional cost. These results confirm and extend those of Experiment 1 in a different task and with different participants, and further suggest that performing confidence estimates during a bimodal task was not more difficult than doing so during the hardest unimodal task (in this case, auditory), despite it requiring the computation of confidence across two perceptual domains. We take this as evidence in support of domain-generality in perceptual metacognition. Second, we found a positive and significant correlation in metacognitive efficiency between the auditory and audiovisual conditions, and a trend between the visual and audiovisual conditions, later replicated in Experiment 3. As in Experiment 1, these results cannot be explained by confounding correlations with first-order performance. We take this as another indication that common mechanisms underlie confidence computations for perceptual tasks on unimodal and bimodal stimuli. While the reported correlations involved a rather low number of participants and were arguably sensitive to outliers (McCurdy et al., 2013), we note that they were replicated several times, under different conditions and tasks in different groups of participants, which is likely in less than 1% of cases under the null hypothesis (binomial test).

The next piece of evidence we brought in favor of domain-general metacognition goes beyond correlational evidence, and provides new insights regarding the mechanisms involved in confidence estimates when the signal extends across two sensory modalities. Using a modeling approach, we found that data in the audiovisual condition was better predicted by models that computed confidence with a supramodal format, based on the information from both the auditory and visual modalities, as compared to simpler models that considered only either one of the two modalities. We take this as evidence in favor of the first mechanism we introduced, according to which metacognition is domain-general because monitoring operates on supramodal confidence estimates, computed with an identical neural code across different sensory modalities. Importantly, while our modeling analyses argue clearly against single domain models as predictors of participants’ behaviour, they could not clearly distinguish between the domain-general average model and the single-modality minimum confidence model. In Experiment 2, the differences in intensity between the left and right stimuli of the auditory and visual pairs were yoked: the staircase procedure we used controlled both pairs simultaneously, increasing (decreasing) the difference between the left and right stimuli in both modalities after an incorrect (two correct) response. As a result, we sampled values from a single diagonal in the space of stimulus intensities, which limits the modeling results. In future studies, non-yoked stimuli pairs could be used—albeit at the cost of a longer experimental session— to explore wider sections of the landscape of confidence as a function of internal signal to better test the likelihood of the models studied here.

Finally, we assessed in Experiment 3 whether domain-general metacognition could arise due to the second mechanism we introduced, according to which domain-generality is driven by the influence of non-perceptual, decisional signals during the computation of confidence estimates. For this purpose, we replicated correlations in metacognitive efficiency between the visual and audiovisual conditions, while examining the neural mechanisms of visual and audiovisual metacognition preceding the perceptual judgment (i.e., at a decisional level). In a response-locked analysis with confidence and condition as within-subject factors, we found that confidence preceding the type 1 response was reflected in ERP amplitude and alpha power (main effect), within a region of interest that included the parietal and sensorimotor cortex corresponding to the hand used for the type 1 task, as well as more frontal sites. Before discussing the main effects of confidence, we note that the analysis also revealed interactions between confidence and condition, revealing that idiosyncratic mechanisms are also at play during the metacognitive monitoring of visual vs. audiovisual signals, and that modulations of ERP and alpha power as a function of confidence were overall greater in the visual vs. audiovisual condition. Regarding the main effects, we found an inversion ERP polarity over left sensorimotor regions, suggesting a link between confidence and selective response activation, so that trials with high confidence in a correct response were associated with stronger motor preparation (Eimer, 1998; Bujan et al., 2009). Regarding alpha power, we found relative alpha desynchronization in occipito-parietal regions, which has been shown to reflect the level of cortical activity, and is held to correlate with processing enhancement (Pfurtscheller, 1992). At the cognitive level, alpha suppression is thought to instantiate attentional gating, so that distracting information is suppressed (i.e. Foxe & Snyder, 2011; Pfurtscheller & Da Silva, 1999; Klimesch, 2012). Indeed, higher alpha power has been shown in cortical areas responsible for processing potentially distracting information, both in the visual and audiovisual modalities (Foxe, Simpson, & Ahlfors, 1998). More recently, pre-stimulus alpha power over sensorimotor areas was found to be negatively correlated with confidence (Baumgarten, Schnitzler, & Lange, 2014; Samaha, Iemi, Postle, 2016), or attentional ratings during tactile discrimination (Whitmarsh, Oostenveld, Almeida, & Lundqvist, 2016). Although these effects are usually observed prior to the onset of an anticipated stimulus, we observed them prior to the type 1 response, suggesting that low confidence in correct responses could be due to the effect of inattention to common properties of first-order task execution such as motor preparation or reaction time (stimulus locked-analyses that are not reported here revealed no effect of confidence prior to stimulus onset). This is compatible with a recent study showing that transcranial magnetic stimulation over the premotor cortex before or after a visual first-order task disrupts subsequent confidence judgments (Fleming et al., 2015).

The finding of lower alpha power with confidence in correct responses is compatible with the observation that participants with more distinct reaction times between correct and incorrect responses had better metacognitive efficiency, as revealed by the correlation between confidence and reaction times overlaps following correct vs. incorrect responses. Thus, attention to motor task execution may feed into the computation of confidence estimates, in a way that is independent of the sensory modality involved, thereby providing a potential decisional mechanism for domain-general metacognition. In experiment 1 we also found that confidence and reaction times overlap were more correlated in the visual condition compared to the tactile, auditory, or audiovisual conditions. Based on these results, we speculate that decisional parameters in link with processes related to movement preparation inform metacognitive monitoring. Our EEG results and the correlations between reaction time and confidence overlaps suggest that decisional parameters may have a stronger weight in the visual than in the other modalities, which could explain the relative superiority of visual metacognition over other senses. We argue that this decisional mechanism in metacognition is compatible with the domain-generality hypothesis, in addition to the supramodal computation of confidence supported by our behavioral and modeling results.

## Conclusion

Altogether, our results highlight two non-mutually exclusive mechanisms for the finding of correlated metacognitive efficiencies across auditory, tactile, visual and audiovisual domains. First, our modeling work showed that confidence estimates during an audiovisual congruency task have a supramodal format, following computations on the joint distribution or on the comparisons of the auditory and visual signals. Thus, metacognition may be domain-general because of supramodal formats of confidence estimates. Second, our electrophysiological results revealed that increased confidence in a visual or audiovisual task coincided with the amplitude of ERP and decreased alpha power prior to type 1 response, suggesting that decisional cues may be a determinant of metacognitive monitoring. Thus, metacognition may be domain-general not only because confidence estimates are supramodal by nature, but also because they may be informed by decisional and movement preparatory signals that are shared across modalities.

## Methods

### Participants

A total of 50 participants (Experiment 1: 15 including 8 females, mean age = 23.2 years, SD = 8.3 years; Experiment 2: 15 including 5 females, mean age = 21.3 years, SD = 2.6 years; Experiment 3: 20 including 6 females, mean age = 24.6 years, SD = 4.3 years) from the student population at the Swiss Federal Institute of Technology (EPFL) took part in this study, in exchange for monetary compensation (20 CHF per hour). All participants were right-handed, had normal hearing and normal or corrected-to-normal vision, and no psychiatric or neurological history. They were naive to the purpose of the study and gave informed consent, in accordance with institutional guidelines and the Declaration of Helsinki. The data from two participants were not analyzed (one in Experiment 1 as the participant could not perform the auditory task, and one from Experiment 2 due to a technical issue with the tactile device).

### General procedure

All three experiments were divided into two main phases. The first phase aimed at defining the participant’s threshold during a perceptual task using a 1-up/2-down staircase procedure (Levitt, 1971). In Experiment 1, participants indicated which of two stimuli presented to the right or left ear (auditory condition), wrist (tactile condition), or visual field (visual condition) was the most salient. Saliency corresponded respectively to auditory loudness, tactile force, and visual contrast (see below for details). In Experiment 2, participants indicated whether the two most salient stimuli among two simultaneous pairs were presented to the same or different ear (auditory condition), visual field (visual condition), or whether the side of the most salient auditory stimulus corresponded to the side of the most salient visual one (audiovisual condition). Stimuli were presented simultaneously for 250 ms. All staircases included a total of 80 trials and lasted 5 min. All thresholds were defined as the average stimulus intensity during the last 25 trials of the staircase procedure. All staircases were visually inspected, and restarted in case no convergence occurred by the end of the 80 trials (*i.e.*, succession of multiple up/down reversals). The initial stimulation parameters in the audiovisual condition of Experiments 2 and 3 were determined by a unimodal staircase procedure, applied successively to the auditory and visual condition.

In the second phase, participants did the same perceptual task, starting with the stimulus intensity determined in phase 1. Another 1-up/2-down staircase procedure was used to keep task performance around 71% throughout phase 2, taking into account training or fatigue effects. Immediately after providing their response on the perceptual task, participants reported their confidence on their preceding response on a visual analog scale using a mouse with their right hand. The left and right end of the scale were labeled “Very unsure” and “Very sure”, and participants were asked to report their confidence as precisely as possible, trying to use the whole scale range, and validate their response with a left click. During a training phase of 10 trials, the cursor turned green/red upon clicking after a correct/incorrect response on the perceptual task. No feedback was provided after the training phase. To exclude trials with trivial mistakes, participants could use the right click to indicate when they had pressed the wrong button, or other obvious lapses of attention. In the audiovisual condition of Experiments 2 and 3, auditory and visual stimuli intensities were yoked, so that a correct (incorrect) answer on the bimodal stimulus led to an increase (decrease) in the stimulus intensity in both modalities. Each condition included a total of 400 trials, divided into 5 blocks. The three conditions (two in Experiment 3) were run successively in a counterbalanced order. One entire experimental session lasted 3 hours.

### Stimuli

Audiovisual stimuli were prepared and presented using the Psychophysics toolbox (Brainard, 1997; Pelli, 1997; Kleiner, Brainard, Pelli 2007) in Matlab (Mathworks). The auditory stimuli consisted of either a 1100 Hz sinusoidal (high pitch “beep” sound) or 200 Hz sawtooth function (low pitch “buzz” sound), played through headphones in stereo for 250 ms with a sampling rate of 44100 Hz. The loudness between the two ears was manipulated to control for task performance. In phase 1, the initial inter-ear intensity difference was 50%, and increased (decreased) by 1% after each incorrect (two correct) answers. The initial difference and step size were adapted based on individual performance. The initial difference in phase 2 was based on the results from phase 1, and the step size remained constant. In the auditory condition of Experiments 2, both sounds were played simultaneously in both ears, and were distinguished by their timber. When necessary, a correction of hearing imbalance was performed prior to the experiment to avoid response biases.

Tactile stimuli were delivered on the palmar side of each wrist by a custom-made vibratory device, using coin permanent-magnetic motors (9000 rpm maximal rotation speed, 9.8 N bracket deflection strength, 55 Hz maximal vibration frequency, 22 m/s^2^ acceleration, 30 ms delay after current onset) controlled by a Leonardo Arduino board through pulse width modulation. Task difficulty was determined by the difference in current sent to each motor. In phase 1 the initial inter-wrist difference was 40%, and increased (decreased) by 2% after each incorrect (two correct) answers. The initial difference and step size were adapted individually, based on subjects performance. A correction of tactile imbalance due to a difference of pressure between the vibrator and the wrist was performed prior to the experiment to avoid response biases. The initial difference in phase 2 was based the results from phase 1 and the step size remained constant.

Visual stimuli consisted in pairs of two 5° x 5° gabor patches (5 cycles/^o^, 11° center-to-center distance). When only one pair was presented (visual condition of Experiment 1 audiovisual condition of Experiments 2 and 3, requiring a laterality judgment), it was vertically centered on the screen. When two pairs were presented (visual condition of Experiment 2 and 3, requiring a congruency judgment), each pair was presented 5.5° above/below the vertical center of the screen. Visual contrast was manipulated, starting with a difference of contrast between gabor patches of 40%, and an increment (decrement) of 2.5% after one incorrect (two correct) answers.

### Behavioural analysis

The first 50 trials of each condition were excluded from analysis as they contained large variations of perceptual signal. Only trials with reaction times between 100 ms and 3 s for the type 1 task and type 2 task were kept (corresponding to an exclusion of 22.2% of trials in Experiment 1 and 12.6% in Experiment 2). In Experiment 3, we used a more lenient superior cutoff of 5 s, resulting in 3.7 % excluded trials, as many trials had to be removed due to artifacts in the EEG signal. Meta-d’ was computed with the Matlab (Mathworks) toolbox provided by Maniscalco & Lau (2012, 2014), with confidence binned into 6 quantiles per participant and per condition. All other behavioural analyses were performed with R (2016), using notably the afex (Singmall et al., 2015) BayesFactor (Morey & Rouder 2015) and ggplot2 (Wikham, 2009) packages. The overlap between confidence and reaction times probability density functions after correct and incorrect responses was computed with the overlap package (Meredith & Ridout, 2014). In all ANOVAs, degrees of freedom were corrected using the Greenhouse-Geisser method.

### Preprocessing of EEG data

Continuous EEG was acquired at 1024 Hz with a 64-channels Biosemi ActiveTwo system referenced to the common mode sense-driven right leg ground (CMS-DRL). Signal preprocessing was performed using custom Matlab (Mathworks) scripts using functions from the EEGLAB (v 13.5.4, Delorme & Makeig, 2004), Adjust (Mognon, Jovicich, Bruzzone, & Buiatti, 2011) and Sasica toolboxes (Chaumon, Bishop, & Busch, 2015). The signal was first downsampled to 512 Hz and band-pass filtered between 1 and 45 Hz (Hamming windowed sinc finite impulse response filter). Following visual inspection, artifact-contaminated electrodes were removed for each participant, corresponding to 3.4% of total data. Epoching was performed at type 1 response onset. For each epoch, the signal from each electrode was centered to zero and average-referenced. Following visual inspection and rejection of epochs containing artifactual signal (3.9% of total data, SD = 2.2%), independent component analysis (Makeig, Bell, Jung, & Sejnowski, 1996) was applied to individual data sets, followed by a semi-automatic detection of artifactual components based on measures of autocorrelation, correlation with vertical and horizontal EOG electrodes, focal channel topography, and generic discontinuity (Chaumon et al., 2015). Automatic detection was validated by visually inspecting the first 15 component scalp map and power spectra. After artifacts rejection, epochs with amplitude changes of ±100 μν DC-offset were excluded (2.9 % of epochs, SD = 3.1%), and the artifact-contaminated electrodes were interpolated using spherical splines (Perrin, Pernier, Bertrand, & Echallier, 1989).

### Statistical analyses of EEG data

Analyses were performed using custom Matlab scripts using functions from the EEGLAB (Delorme & Makeig, 2004) and Fieldtrip toolboxes (Oostenveld, Fries, Maris, & Schoffelen, 2011). Event-related potentials were centered on zero. Time-frequency analysis was performed using Morlet wavelets (3 cycles) focusing on the 8-12 Hz band. Voltage amplitude and alpha power were averaged within 50 ms time windows, and analyzed with linear mixed effects models using R together with the lme4 and lmerTest packages (Bates, Maechler, Bolker, & Walker, 2014; Kuznetsova, Brockhoff, & Christensen, 2014). Models were performed on each latency and electrode for individual trials, including raw confidence rating and condition (i.e., visual vs. audiovisual) as fixed effects, and random intercepts for subjects. Random slopes were not added as they induced convergence failures (i.e., parsimonious instead of maximal models, see Bates et al., 2015). Statistical significance for ERPs and alpha power within the region of interest was assessed after correction for false-discovery rate. Topographic analyses were exploratory, and significance was considered for p < 0.001 without correcting for multiple comparisons.

### Modeling Procedure

Our models assume that confidence in each trial is proportional to the likelihood of a correct answer, given a joint distribution of the internal signal associated with the two pairs of visual or auditory stimuli. Relying on two parameters, we aimed at calculating the proportion of trials in which confidence was higher/lower than the median confidence value, for correct/incorrect type 1 response, for congruent/incongruent stimuli. For each participant, we estimated the internal noise (σ) and confidence criterion (*c*) that best fitted the proportion of trials in the 8 corresponding categories (2 confidence bins x 2 accuracies x 2 conditions).

### Model of one participant

We modeled each participant’s internal signal as a bivariate normal. Each dimension corresponded to one of the stimuli pairs in each condition. The bivariate distribution was parametrically defined with an arbitrary mean of μ = (1,1) and two standard deviations σ1, σ2. In the bimodal condition, σ1 and σ2 corresponded to the internal noise for the visual and auditory signal respectively, and were allowed to vary independently. In the unimodal conditions instead, σ1 and σ2 corresponded to each of the stimuli pairs of the same modality and were therefore constrained to be equal. An additional parameter c determined the criterion above which a decision was associated with high confidence ratings (i.e., type 2 criterion). Thus, the model relied on three assumptions: first, it assumed equal priors for all possible stimuli. Second, type-1 decisions were assumed to be unbiased and optimal. Third, confidence was defined as proportional to the density P(correct|x_1_, x_2_), where x_1_, x_2_ correspond to the strength of the evidence of each pair of stimuli in a given trial. (In models of unimodal conditions, x_1_, x_2_ corresponded to two stimuli pairs in the same modality; whereas in the bimodal conditions they corresponded to the auditory and visual stimuli. We use the notation x_1_, x_2_ for generality). We argue that the assumption of equality for σ_1_ and σ_2_ is a reasonable one in the unimodal visual case, where the two stimuli pairs differed only on their vertical position (but did not differ in their distance from the vertical midline). This assumption however is less clearly valid in the unimodal auditory condition, where the two pairs of stimuli were phenomenologically different (a sinewave ‘beep’ vs. a sawtooth ‘buzz’). We note that the model was flexible enough to fit the different behavioural patterns of most participants, and that the model fits obtained for the unimodal auditory condition were comparable to those in the unimodal visual condition (see Results).

### Modeling strategy

We first estimated the σ and *c* parameters from the unimodal data for each participant, and then combined them under different models to estimate their fits to the audiovisual data (See Figure S2). Note that with this procedure, and unlike the fits to the unimodal conditions, the data used to estimate the model parameters were different from those on which the model fits were compared. We grouped the models into three families to compare them systematically. The family of *domain-general models* echoes the unimodal models and represent the highest degree of integration: here, confidence is computed on the basis of the joint distribution of the auditory and visual modalities (Figure 4a). Within this family, the *average model* considers one value of σ for each modality and takes a criterion resulting from the mean of the two modalities estimated. The family of *comparative models* (Figure 4b) assumes that confidence can only be computed separately for each modality and combined into a single summary measure in a second step. Within this family, the *minimum-confidence model* takes the minimum of the two independent confidence estimates as a summary statistic. Finally, the family of Single-modality models (figure 4c), assumes that confidence varies with the internal signal strength of a single modality and therefore supposes no integration of information at the second-order level. Within this family, the *maximum efficiency model* computes confidence on the basis of the modality with the best metacognitive efficiency alone. We calculated the Bayesian information criterion (BIC) to compare the different models while accounting for differences in their number of parameters.

Our modeling approach can be seen as an extension of recent work (Aitchinson Bang, Bahrami, & Latham, 2015), where two-dimensional SDT models similar to the ones we developed here revealed that participants estimated their confidence as the likelihood of a correct response given the sensory data. These models were defined for a 2-interval forced-choice task, where the two intervals were mutually exclusive and the signal strength in one interval carried information about the alternative interval. Consequently, confidence increased with increasing signal strength in one interval, and *decreasing* signal strength in the other interval. In our task, on the other hand, congruency judgments had to be made on the basis of both stimulus pairs; and the strength of the sensory signal in one modality did not carry information about the second modality. Thus, in our models, the estimated confidence increased with increasing signal strength in both dimensions considered (see Figure S2, cf. Figure 9 from Aitchinson et al).

### Decision rule - type 1 task

We modeled the type 1 decision (*i.e.*, the congruency judgment) as a ratio of log likelihoods.

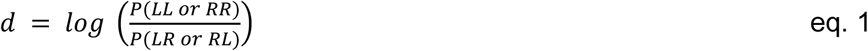

Where LL, RR, LR and RL correspond to the four possible stimuli combinations of stimuli x_1_, x_2_ most salient on the left (L) or right (R) side. *LL* and *RR* correspond to congruent stimuli pairs and *LR, RL* to incongruent stimuli pairs. It follows from eq. 1 (for the derivations, see SI) that the optimal type-1 criteria are placed at x_1_ = 0 and x_2_ = 0.

### Confidence judgment - type 2 task

We modeled the type-2 decision (*i.e.*, the confidence judgment) as the probability of having given a correct response, given the stimulus strength. We then tested eight different models under which confidence may be computed (see SI for schematics representing the predictions and BIC values for all models). In the domain-general models, confidence is computed on the basis of the joint bivariate distribution in the bimodal condition. Thus,

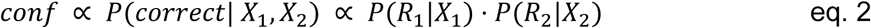

The three members of this family differed in the values of σ used for each of the terms.

In the three domain-specific models considered, confidence is computed independently for each dimension and, in a subsequent step, a summary measure is reported: either the maximum of the two computed confidence values (Maximum confidence eq. 3), the average of the two (Mean confidence, eq. 4) or the minimum of the two (Minimum confidence, eq. 5).

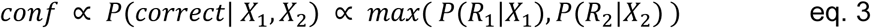

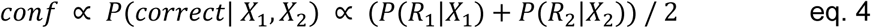

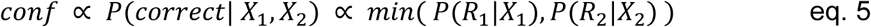

In the single-domain models, confidence is calculated based only on a single modality, namely the one with the highest or lowest metacognitive efficiency as measured in the unimodal conditions (Maximum or Minimum metacognition model respectively, eq. 6).

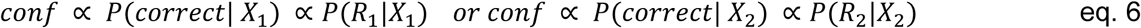

### Model fits

We first estimated the percentage of variance explained by the unimodal models. To calculate R^2^, we first used the *nlme* package in R (Pinheiro et al., 2016) to estimate the predictive power of our models while allowing for random intercepts for each participant. We then used the *piecewiseSEM* package (Lefcheck, 2015) to estimate the percentage of variance explained, following the methods developed by Nagakawa et al. (2013). BIC weights for the model fits to the bimodal condition were estimated following (Burnham & Anderson, 2002) and the details are described elsewhere (Solovey et al., 2014).

## Acknowledgements

NF was an EPFL fellow co-funded by Marie Skłodowska-Curie. O.B. is supported by the Bertarelli Foundation, the Swiss National Science Foundation, and the European Science Foundation. We thank Shruti Nanivadekar and Michael Stettler for their help during data collection.

## Author Contributions

NF and EF developed the study concept and contributed to the study design. Testing and data collection were performed by NF. NF and EF performed the data analysis. EF and GS performed modeling work. NF and EF drafted the paper; all authors provided critical revisions and approved the final version of the paper for submission.

## Supplementary Information

### Supplementary behavioural results

**Experiment 1.** No effect of condition on response criterion [F(1.96,27.47) = 0.30, p = 0.74, η_p_^2^ = 0.02] nor on average confidence was found [F(1.87.26.21) = 2.42. p = 0.11, ηp2 = 0.15]. As reported previously (Ais et al. 2016). average confidence ratings correlated between the auditory and visual conditions [adjusted R^2^ = 0.26. p = 0.03]. between the tactile and visual conditions [adjusted R^2^ = 0.55. p = 0.001]. and between the auditory and tactile conditions [adjusted R^2^ = 0.51. p = 0.002]. No effect of condition on type 1 reaction times [F(1.78.24.96) = 0.28. p = 0.73, η_p_^2^ = 0.02] or type 2 reaction times [F(1.77.24.84) = 1.77. p = 0.39, η_p_^2^ = 0.06]. was found.

**Experiment 2.** As in Experiment 1. no effect of condition on response criterion [F(1.87.24.27) = 2.12, p = 0.14, η_p_^2^ = 0.14] nor on average confidence was found [F(1.76.24.64) = 0.91. p = 0.40. η_p_^2^ = 0.06]. No evidence of multisensory integration was found at the first-order level. as the perceptual thresholds determined by the staircase procedure were not lower in the bimodal vs. unimodal conditions [p = 0.17]. This is likely due to the task at hand involving a congruency judgment. Average confidence ratings correlated between the auditory and audiovisual conditions [adjusted R^2^ = 0.56. p = 0.001]. between the visual and audiovisual conditions [adjusted R^2^ = 0.38. p = 0.01]. and a trend was found between the auditory and visual conditions [adjusted R^2^ = 0.12. p = 0.11]. A significant main effect of condition on type 1 reaction times [F(1.66.21.53) = 18.05. p < 0.001, η_p_^2^ = 0.58] revealed faster responses in the visual [1.30 s ± 0.10 s] compared to the auditory [1.47 s ± 0.13 s] and audiovisual task [1.68 s ± 0.11 s]. No difference was found for type 2 reaction times [F(1.82.23.62) = 1.69. p = 0.21, η_p_^2^ = 0.11].

**Experiment 3.** Contrary to what was found in Experiments 1 and 2. response criterion varied across conditions [t(17) = 4.33. p < 0.001. d = 0.63]. with a tendency to respond "congruent” more pronounced in the audiovisual [mean criterion = 0.27 ± 0.12] vs. visual condition [mean criterion = -0.02 ± 0.15]. This effect was unexpected but did not preclude from running subsequent analyses dealing with metacognitive sensitivity that are independent of response criterion. We found no effect on average confidence [t(17) = 0.56. p = 0.14. d = 0.08]. Average confidence ratings correlated between the visual and audiovisual conditions [adjusted R^2^ = 0.65. p < 0.001].

**Figure S1:**
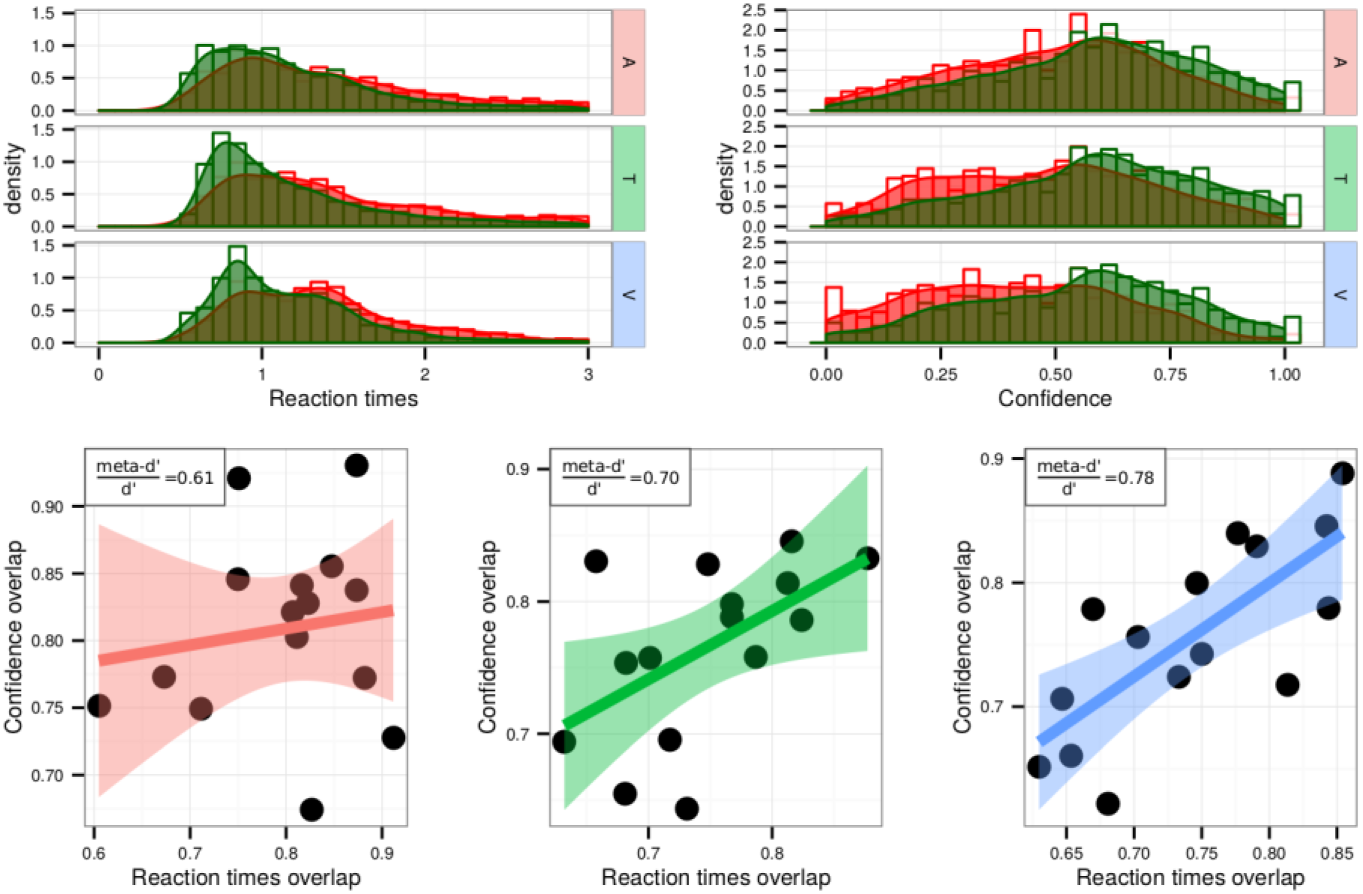
Reaction times and confidence overlap. **Upper row**. Histogram and density functions of reaction times (left) and confidence ratings (right) following correct (in green) and incorrect type 1 responses (in red), for a representative participant in the auditory (A), tactile (T), and visual (V) conditions. **Lower row.** Correlation between the reaction times and confidence overlaps following correct and incorrect type 1 responses across participants. The metacognitive efficiency (meta-d’/d’) for each modality is mentioned on the top-right corner of each plot. Note that the correlation strength varies accordingly to metacognitive efficiency. The auditory condition is represented in red, the tactile condition in green, and the visual condition in blue.

### Modeling

As described in the Methods, LL, RR, LR and RL correspond to the four possible stimuli combinations of the internal signal intensity x_1_, x_2_, with LL and RR corresponding to congruent stimuli pairs and LR, RL corresponding to incongruent stimuli pairs.

#### Type-1 decision for all models

According to the type-1 decision rule, the decision will be “congruent” if *d > 0* and “incongruent” if *d < 0,* where:

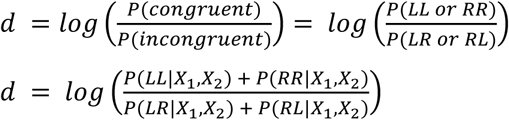

Applying Bayes’ rule and assuming equal priors:

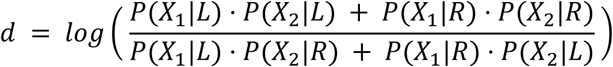

And it follows that the criterion for *d = 0* where the congruent and incongruent stimuli are equally likely, should satisfy the relation:

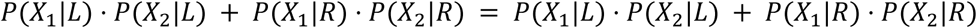

The two solutions to this relation are given by

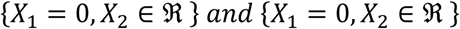

#### Type-2 decision (confidence judgment)

We considered eight different ways in which confidence could be computed in the bimodal audiovisual condition.

Under the domain-general models:

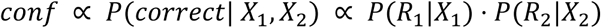

Assuming bivariate normal distributions of the internal signals, and that each modality takes the σ value it can be shown that

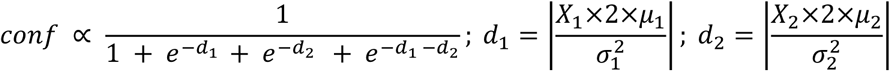

Under the domain-specific. minimum confidence model:

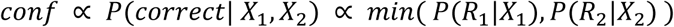

And hence:

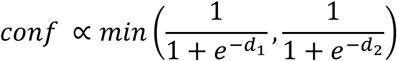

And finally. under the second domain-specific. mean confidence model:

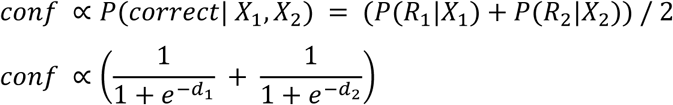

#### Individual model fits

**Figure S2:**
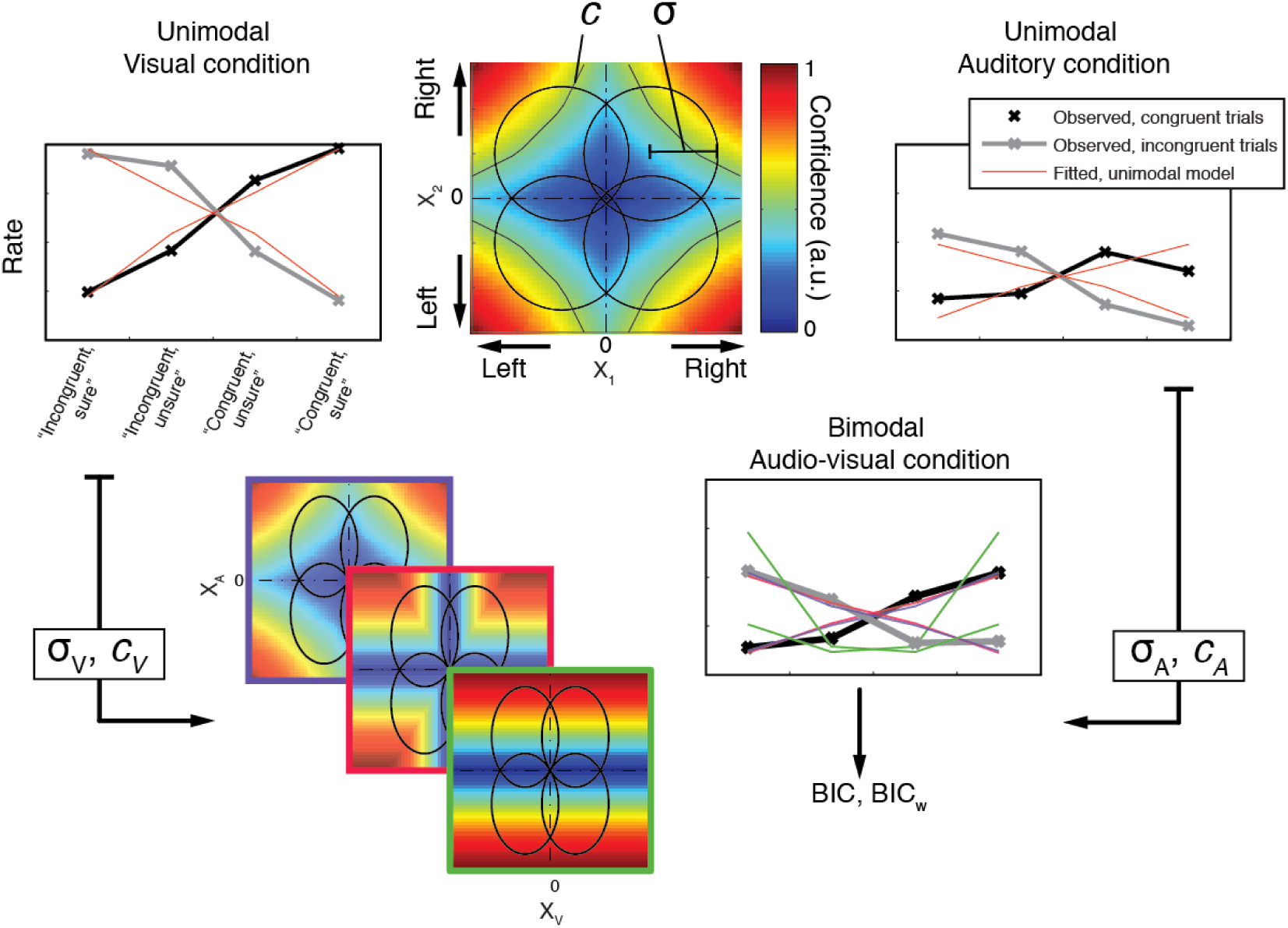
Modeling strategy. We modeled the unimodal data (top panel) to obtain estimates of the modality-specific internal noise (σ) and criterion (c) for each participant. We then combined these values to predict the bimodal data of the audiovisual condition. according to three rules depicted by colors (bottom panel). Blue stands for domain-general models. red stands for comparative models. and green stands for single-modality models. We estimated BIC and BICw to compare the candidate models in the bimodal conditions.

**Figure S3:**
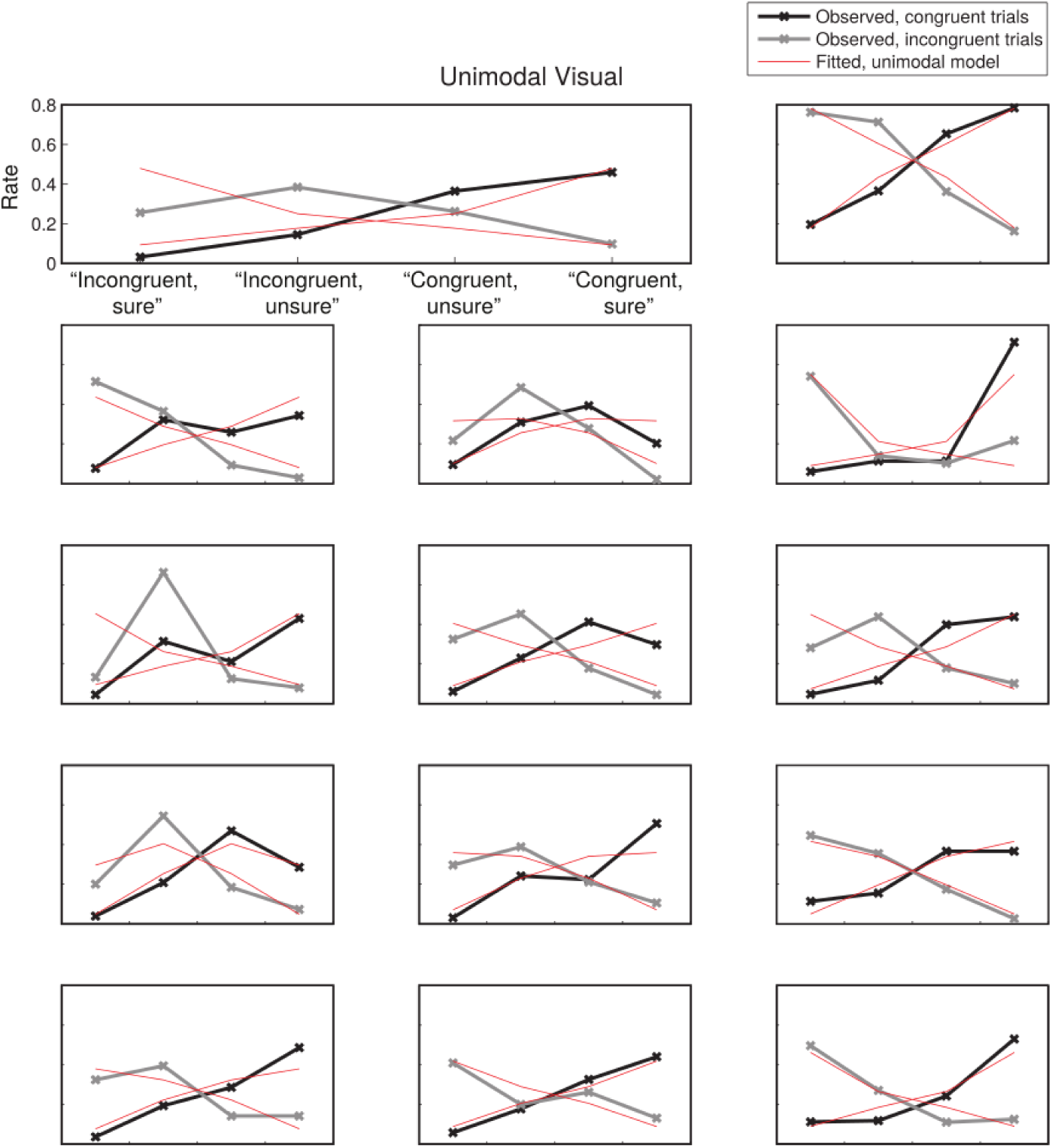
Individual model fits for the unimodal visual condition. Each plot shows the response rates for a participant in experiment 2. Two kinds of trials were presented: congruent (represented with black lines) and incongruent (represented with grey lines). After a median split of each participant’s confidence ratings, four possible responses resulted from the combination of the congruency judgment (“congruent”/”incongruent”) and the confidence judgment (“high confidence”/”low confidence”, represented here as “sure”/”unsure”). The red lines represent the response rates predicted by the unimodal model. These model predictions are presented here for illustration purposes only. They are the result of searching the parameter space (σ, c) for the values that best predicted the 8 response rates. While they suggest that the unimodal model we used was plausible, they were not subjected to any model comparison.?

**Figure S4:**
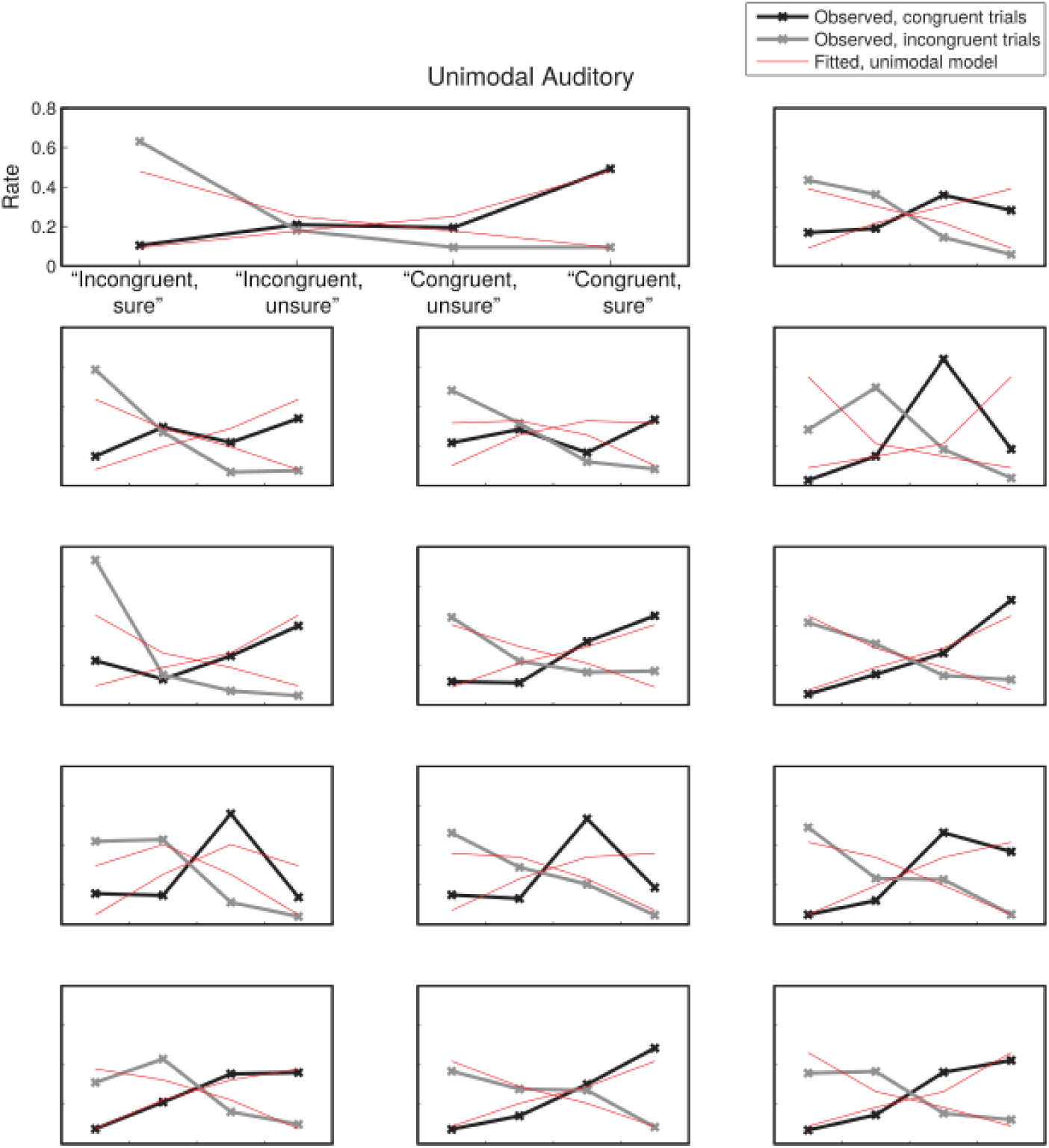
Individual model fits for the unimodal auditory condition. See legend in figure S2 for details.

**Figure S5:**
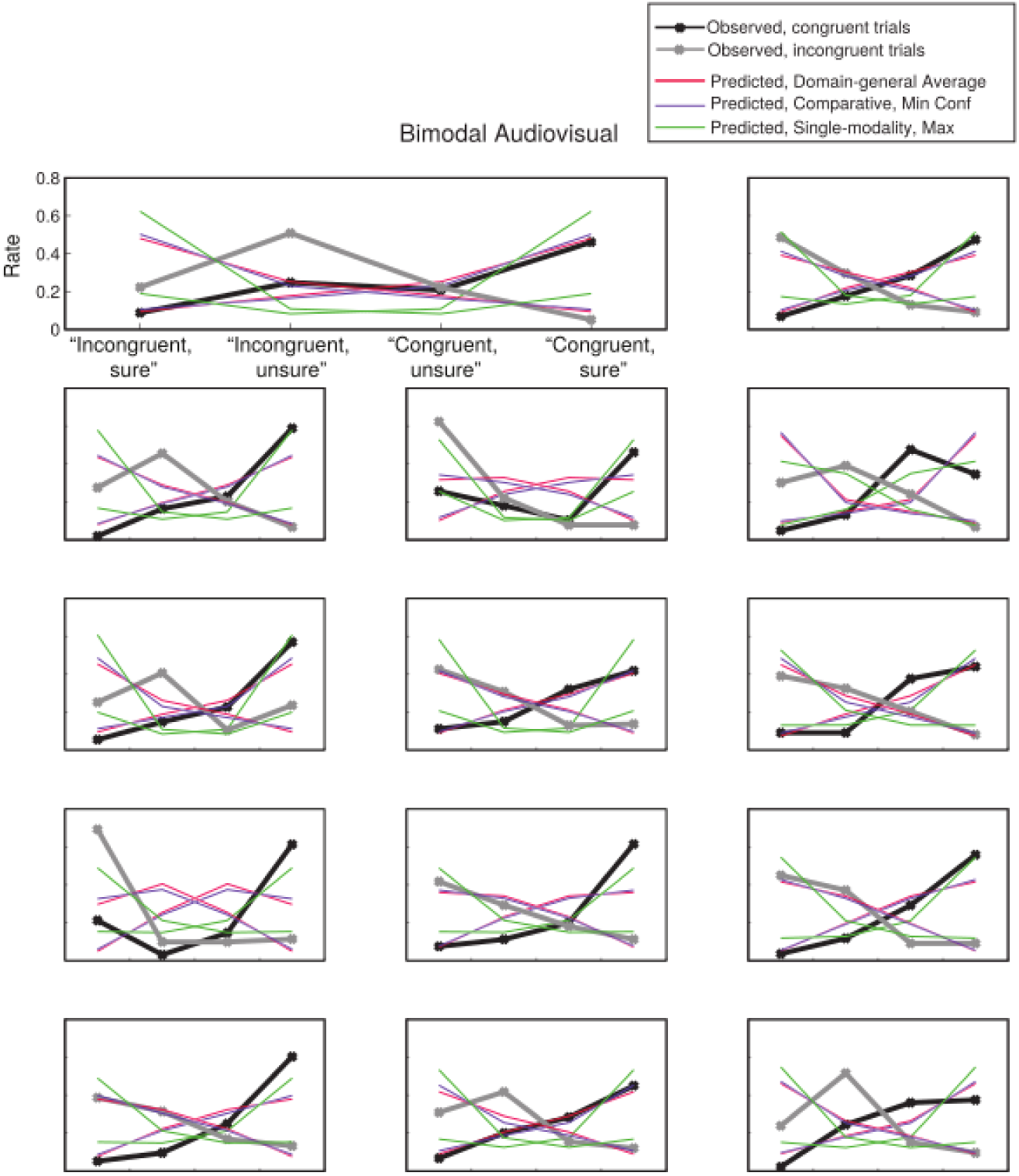
Observed and predicted response rates for each participant. for the three best models (namely. the domain-general Mean model. the domain-specific conservative Minimum model and the single-domain Maximum metacognition model). Each model predicted response rates for 8 categories that result from combining the three factors: stimulus congruency (congruent/incongruent). response (correct/incorrect) and binned confidence (high/low). High and low confidence responses were defined as above or below each participant’s median confidence, across all conditions.

**Figure S6:**
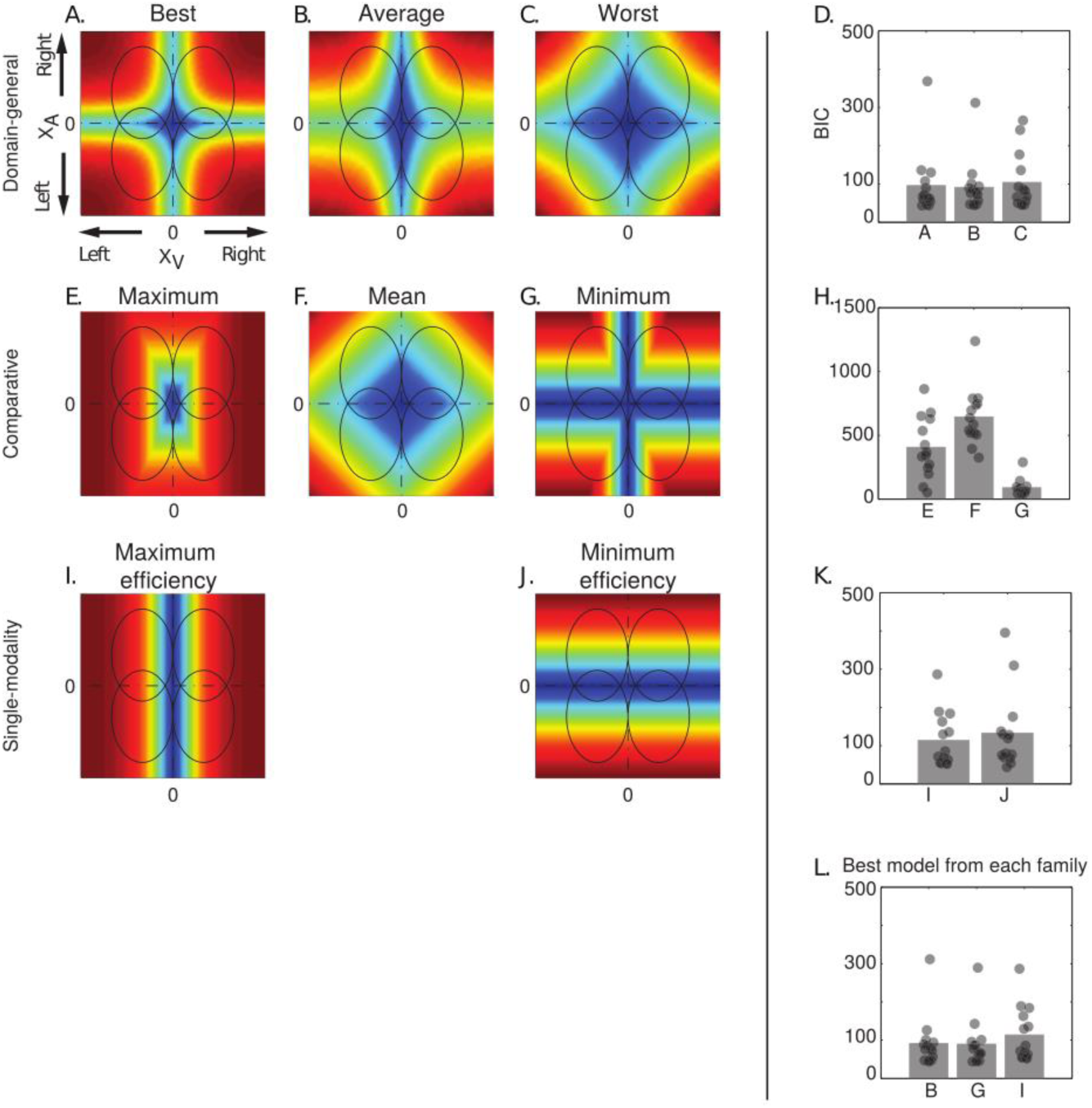
Left panel: confidence model predictions for all models tested. Ellipses represent the partially overlapping bivariate internal signal distributions for each of the stimuli combinations, represented at a fixed density contour. The models allowed the auditory and visual modalities to have different σ parameters, leading to ellipses instead of circles. Within each schematic of model predictions, the top right quadrant corresponds to congruent stimuli, where the stimuli of both pairs were stronger on the right side. The colors represent the predicted relative confidence for every combination of internal signal strength for each stimulus pair (XA, XV). Note that in all models except the two single-modality ones, confidence increases with increasing evidence in all four quadrants, because the congruency judgment is based on the conjunction of stimuli, and cannot be performed on basis of one stimulus only (see Aitchison et al., 2015 for similar work) **Right panel**: BIC values for the model fits in the audiovisual condition.

